# Kinematic and biomechanical analyses in Drosophila suggests that most legged locomotion in insects can be understood within a single framework

**DOI:** 10.1101/455246

**Authors:** Chanwoo Chun, Tirthabir Biswas, Vikas Bhandawat

**Affiliations:** Department of Biology, Duke University.; Department of Physics, Loyola University, New Orleans.; Duke Institute for Brain Sciences, Duke University.

## Abstract

Despite the fact that all insects have six legs, they display considerable differences in their legged locomotion. Are these differences irreconcilable, or simply different domains of the same system? In this study, we investigate walking in *Drosophila*. We find that despite the fact that flies walk extremely slowly relative to their size, they predominantly employ an alternating tripod gait, a gait typically associated with high-speed locomotion. The kinematics of their center of mass (CoM) is diametrically opposite to the CoM kinematics observed in insect runners such as cockroach. We resolve this tension between similar gaits, and differing kinematics in slow and high-speed locomotion by showing that the mechanics of a tripod gait naturally reduces to a simple biomechanical model which can support a range of kinematic patterns including both fly-like and cockroach-like patterns. These findings suggest that legged locomotion in different insects might represent different domains of the same system.

---

Insects have considerable flexibility in how they employ their six legs. This flexibility is, in part, because access to six legs simplifies stability considerations: As long as insects have three legs on the ground to form a tripodal base of support, they have a considerable discretion in how they choose to employ the other legs. Given this flexibility, any assertion related to insect locomotion is likely to have exceptions. Still it is important to create a general analytical framework to rigorously assess what aspects of an insect’s locomotion can be understood within a single framework, and when observations require fundamentally different framework.

Consider one crucial aspect of legged locomotion – the nature of coordination among legs during walking – or gait. There is a general agreement that insects predominantly employ a tripod gait (see Figure 1 and S1) when they move at high speeds ^1^^-^^7^. However, there is little agreement regarding the coordination pattern between legs at lower speeds: some researchers have proposed that a tripod coordination pattern persists across all speeds ^8,9^. Others have proposed a tetrapod gait (4 legs on the ground, see Supplementary Figure 2) at intermediate speeds, and a metachronal gait at lower speeds ^2-4,10,11^. Moreover, unlike gait transitions in bipeds and quadrupeds that occur at precise speeds, gait transitions in insects are probabilistic. Another related idea is that gaits themselves are not discrete but form a continuum, which has led to the idea that insects employ a continuum of gaits ^8,10,12-16^. Do these different viewpoints result from fundamental differences between insects, idiosyncrasies in data collection and curation, or analytical methods employed to define gait? A rigorous analytical approach is necessary to distinguish between these possibilities.

**Figure 1.**
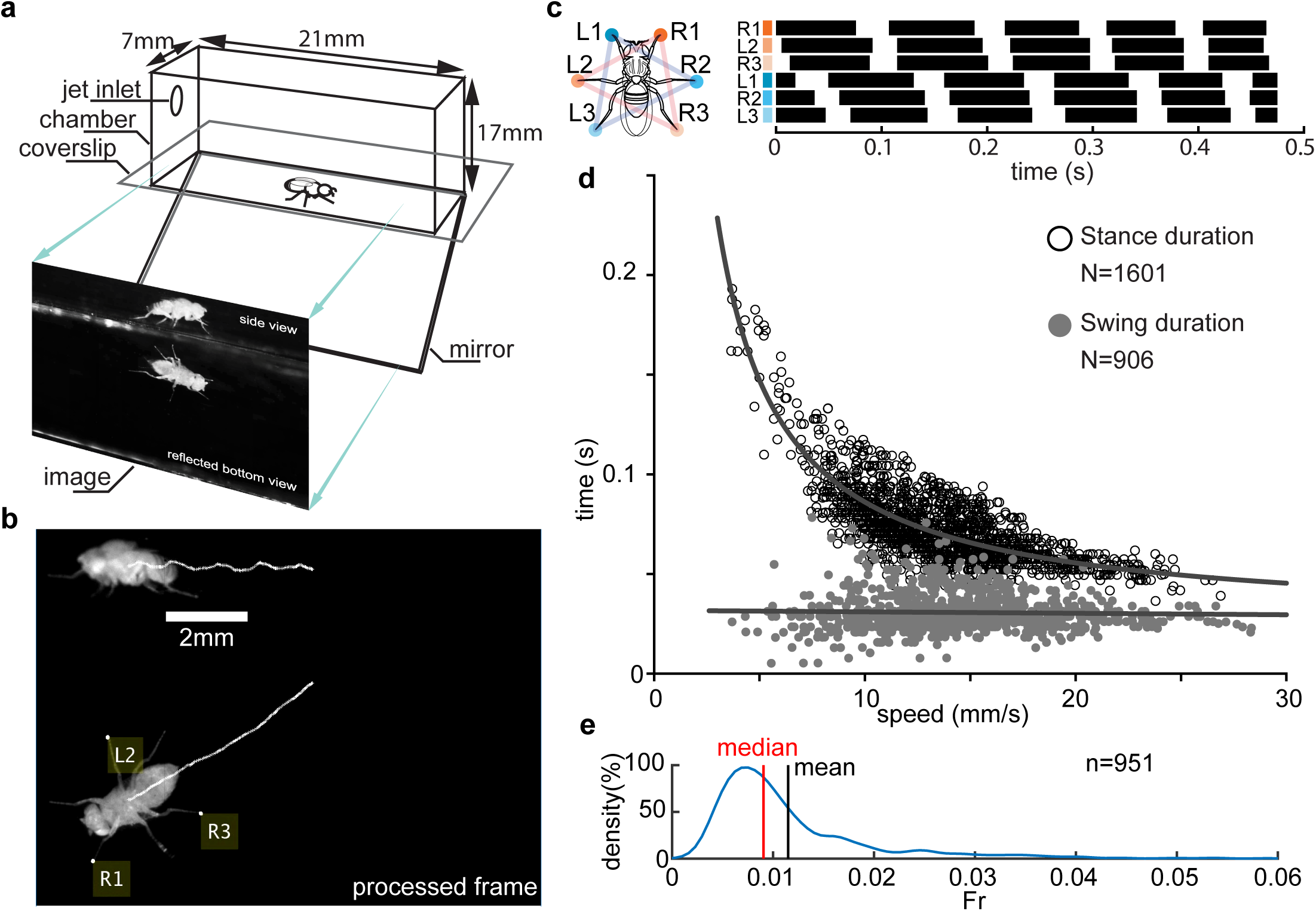
Experimental setup. (**a**) Schematic of the arena. (**b**) Example of a processed video. The white traces track a point on the thorax and is a proxy for the CoM. Yellow labels denote feet that are in stance or the footholds. (**c**) Leg numbering and color schemes as well as the gait maps: right tripod legs in orange; left tripod in blue. Example footfall patterns. Each row is a single leg. Black bar represent stance phases. (**d**) Stance and swing durations plotted against speed. The top dark, grey line is a best fit of reciprocal function to stance durations. The bottom, lighter grey line is a best fit of linear regression to swing durations. (**e**) Distribution of Froude number (*Fr*), a measure of non-dimensional speed obtained using Kernel density estimation show that flies walk unusually slowly. The vertical line represents mean of average Fr.

A similar diversity of ideas can be found in the context of kinematics and kinetics of the center of mass (CoM) during locomotion. Much of the work on understanding CoM kinematics and kinetics has focused on cockroaches while they are running at high speeds^17,18^. An elegant series of studies on running cockroaches has shown a striking similarity to mammalian running – in both cases the CoM reaches a minima in speed and height at mid-stance^17,18^. This kinematic pattern can qualitatively be described as if the CoM is bouncing on a spring ^19-23^. Essentially, a cockroach that is running fast uses an alternating tripod to do so; the three legs of a tripod can be replaced by a single spring-loaded leg which is compressed as it moves forward during the first half of stance, and extends during the second half; the energy stored during the first half of the stance propels the COM during the second half of the stance. In part, because of uncertainty in gait, it has been difficult to perform a similar analysis in the context of most other insects. However, it is clear that the CoM kinematics of some other insects, including stick insects and fruit flies, is fundamentally inconsistent with a cockroach-like spring-mass model because their horizontal speed during walking is at its maximum at mid-stance ^8^^,^^10^. Once again, it is important to understand whether these different kinematic patterns are simply part of a continuum, or represent fundamentally different mechanics.

In an attempt to arrive a general framework for analyzing gaits and CoM kinematics, we created a new automated method for measuring the movement of a fly’s CoM in all three dimensions, while also tracking the position of the fly’s stance legs. Using this method, we analyzed a fly’s gait over >2000 steps during which the fly is always walking straight. We also created a new method for gait analysis, and using this method, show that flies predominantly employ a tripod gait. The three legs of a tripod act like a single leg whose behavior can be described by a simple extension to the spring-loaded leg proposed in the case of cockroach: this extension can be thought of as an angular spring that modulates the forward movement of the CoM. This new biomechanical model – **a**ngular and **r**adial **s**pring **l**oaded **p**endulum or ARSLIP accurately models a fly’s locomotion, and is able to explain how tripod geometry affects the nature of forces that act on the fly, and ultimately defines its kinematics. Finally, we show that fly walking occupy a small region of the ARSLIP parameter space, but other subspaces within the ARSLIP parameter space can produce most kinematic patterns observed in insects. Together these results support the idea that many insect gaits can be studied within a single analytical framework.

## Results

Because of the flexibility associated with limb kinematics and the fact that flies are small, we reasoned that a large dataset with high spatial resolution would be necessary to resolve questions regarding a fly’s gait and its CoM kinematics. Therefore, we started by creating an automated data acquisition system. Similar to an approach employed previously^4^^,^^24^, we recorded the side and the bottom view (reflected off a mirror) of a fly walking in a clear cuboid chamber that enclosed the fly (Figure 1a). To avoid conflating between gaits used during turning, we extracted all the steps during which a fly walked straight for more than one step; these steps constituted a small proportion of the data (see methods for details). The CoM of the fly, and the leg tips during stance were tracked using a custom algorithm detailed in the methods. Briefly, CoM was extracted using the Kanade-Lucas-Tomasi (KLT)^25^ algorithm and produced very reliable estimates of the CoM. The vertical resolution of our data was 20 µm (see Methods 2.2), and at this resolution, the rhythmic up-and-down movement of the CoM is clearly visible (Figure 1b). The legs were denoted using convention followed in previous studies^26^(Figure 1c), and the gait map (Figure 1c) was plotted such that the legs of one tripod – right prothoracic (R1), left mesothoracic (L2), and right metathoracic (R3) are plotted in consecutive rows (rows marked in orange); and those of the other tripod (L1-R2-L3) are plotted in another set of consecutive rows (marked in blue), making it easy to visualize whether the gait is a tripod.

As a means of corroborating previous findings, we plotted stance and swing duration as a function of speed (Figure 1d). In multiple studies across insects, as seen in Figure 1d, the stance duration is inversely proportional to speed and the swing duration is nearly constant ^3^^,^^8^^-^^10^^,^^27^. The range of speed in our dataset is marginally lower than the range of speed observed in other studies ^4^^,^^8^.

An important feature of a fly’s locomotion is that relative to their size they walk at slow speeds. Froude number (*Fr*)– a dimensionless speed defined as 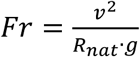 where *v* denotes speed, *R_nat_* is the natural length of a leg, and *g* a gravitational acceleration shows that the *Fr* in our dataset ranges from 0.0012 to 0.059. The median *Fr* is ~0.01; in comparison, a human walking at a leisurely pace of 3 mph walks at a *Fr* of 0.2 or 4.5 times faster in relative terms^28^.

## Flies’ gait is most consistent with an alternating tripod with little evidence for gait transition with speed

A strict definition of gait implies synchronicity between a subset of legs. As an example, a strict definition of tripod implies that the three legs that form the tripod (R1-L2-R3) move synchronously and in an opposite phase to the other tripod (L1-R2-L3). This strict definition is rarely satisfied because of the lack of perfect synchronicity among the legs that are part of a tripod, and because the stance duration is usually longer than the swing duration, it is impossible for the two tripods to be out-of-phase throughout the entire stance. In order to address this problem, consistent with approaches taken by other authors^29^, we will adopt a more flexible definition of gait which allows for small deviations from perfect synchronicity.

As a first step towards assessing gait, we characterized coordination between legs using two methods: First, coordination can be defined based on delays between the start of either the swing or stance phase. Figure 2a shows the relative delays between the times that different legs enter the stance phase. Each row is a single step; and the rows are sorted by speed. The three legs of the tripod enter stance with a short inter-leg delay. Importantly, there is no noticeable change in the coordination pattern as flies walk faster. The raw gait map (Figure 2a) supports a tripod gait over all speeds, this trend can be quantified by calculating the delays relative to the cycle period or the time it takes a leg to complete both a swing and a stance (Figure 2b). The delays normalized to the cycle period are close to zero for legs within a tripod. The prothoracic leg lead the other legs in its tripod with a small, but significant negative delay (Wilcoxin signed-rank test); a similar trend was observed in other studies^2^^,^^26^^,^^30^^,^^31^. On the other hand, the normalized delays for legs across the two tripods is 0.5. Moreover, the median phase does not change significantly with the speed of walking (Wilcoxin rank-sum test).

Second, we employed instantaneous phase to measure coordination between legs (Figure 2c). Instantaneous phase, which has been used to investigate coupling strengths between hexapod legs^29^^,^^32^, is a more accurate measure of coordination since it assesses coordination between legs throughout the step rather than at the beginning of stance (Methods section 3.2). The distribution of instantaneous phase between the reference leg (R1) and the other legs show that the legs that are part of a tripod have a small phase difference (Figure 2D), while legs in the opposing tripod have a large phase difference. The phase plots also show that the front leg of the tripod leads the middle and back legs.

**Figure 2.**
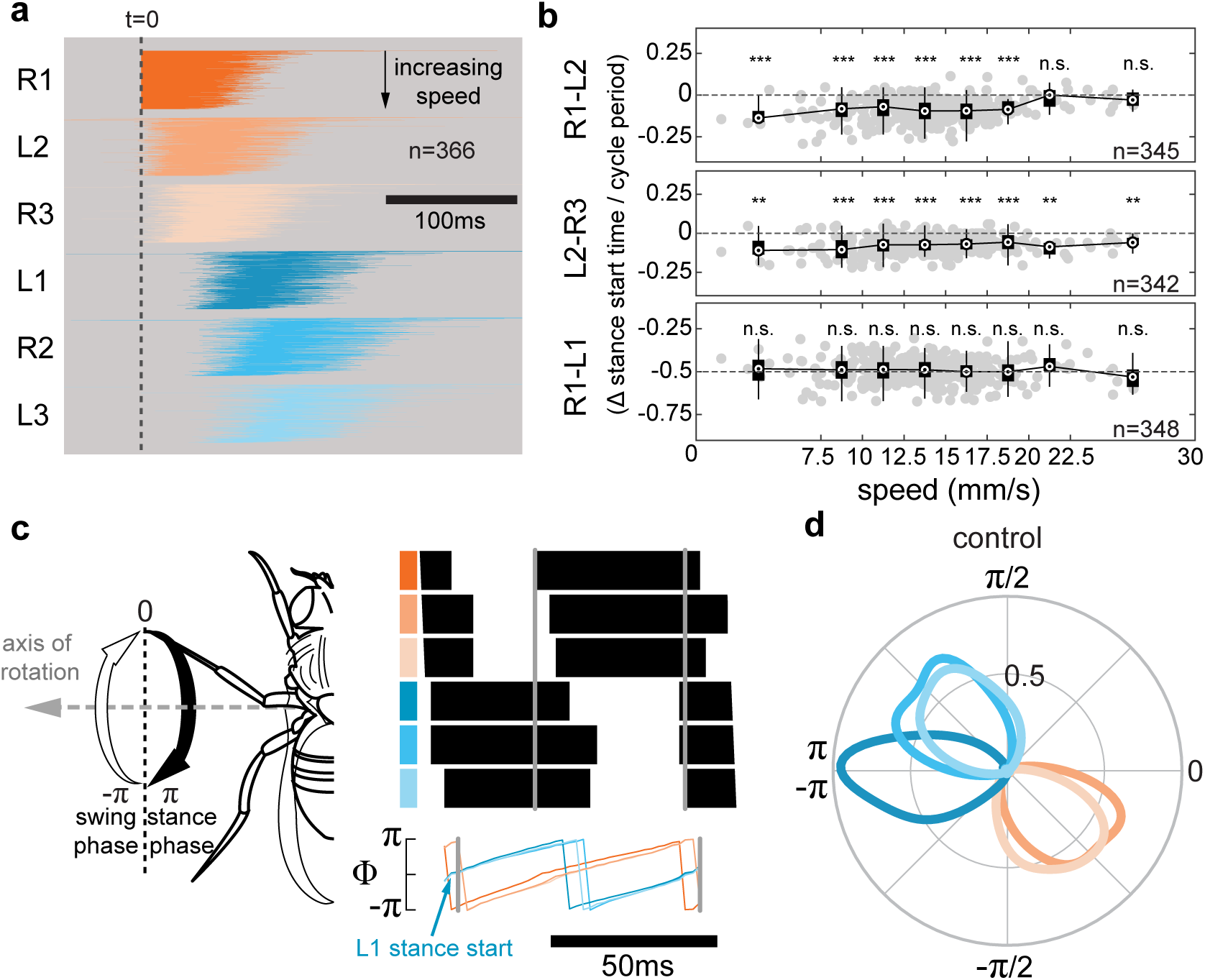
Coordination between legs does not depend on speed and is consistent with a tripod gait. (**a**) Stances phases of all steps relative to R1 sorted by speed. (**b**) Normalized time delays of stance start times between legs within a tripod (R1 and L2, L2 and R3) and legs in the oppositing tripod (R1 and L1). The time delays were normalized by cycle durations. Box plot with edges of the bins as shown on the x-axis. Significance refers to Wilcoxon sign rank test to test that the phase difference is < 0 (for L1-R2 and L2-R3). For R1-L2 (bottom), difference from -0.5 is being tested. (**c**) Definition of leg phase angles. Stance start is at 0, stance end at π; swing start at -π and swing end at 0.(**d**) Histogram of leg phases relative to R1 shows that coordination between leg is consistent with a tripod gait.

The analyses in Figure 2 strongly suggest that the coordination pattern between legs is more consistent with a single gait that is close to the tripod gait. To further assess whether flies predominantly walk using a tripod gait, it is important to summarize the coordination between legs into a single metric. Other studies have attempted to arrive at a single metric which quantifies how close to tripod a given step is. One metric is the tripod index which quantifies how tripodal a particular step is based on the fraction of time only 3 legs are on the ground ^8^. This metric is flawed and largely measures duty factor, or the percentage of the cycle period spent in stance (Supplementary Figure 1). Using tripod index, even a perfect tripod gait would appear to be a mixture of tripod and tetrapod (Supplementary Figure 1). Therefore, we designed a class of metrics – ***gait delay indices (GDIs)*** - that are independent of duty factor. The central idea behind GDIs is that any gait is defined by a fixed pattern of relative delays or phase difference between legs; a unit vector constructed using these delays would always point in the same direction irrespective of the duty factor (Supplementary Figure 1). Therefore, the projection of the empirical delay vector along the direction of the idealized gait – a GDI – is a useful measure of gait (Supplementary Figure 1).

Because there are 6 legs, a set of 5 delays define a gait. GDIs can take multiple forms depending on which of the 5 delays are employed to construct the delay vector. Since, some previous studies have shown that flies transition from tetrapod to tripod ^4^^,^^8^ as the speed of locomotion increases, we designed a GDI (Figure 3a) to distinguish between the tripod and tetrapod gaits. The set of delays that best distinguished tripod and tetrapod are L1-R3, L3-R1, L3-L1 (see Supplementary Figure 2 for the reasoning). Using this set of delays, the projection of an ideal tripod along the two ideal tetrapod directions are 0.6 and 0.3, making it an effective metric for distinguishing between the tripod and tetrapod gaits. GDIs based on this set of delays show a single cluster close to 1, implying that the gait is predominantly a tripod (Figure 3a top). One reason that the GDIs are not centered on 1 is because 1 is the upper bound for GDIs because it is based on projections of unit vector. Another reason is that the gait is not a classical tripod, but a modified tripod (metachronal tripod or m-tripod, see Supplementary Figure 2 for definition) in which L1 leads R2 which leads L3 by a small delay. Almost all the data points are bounded by the pure tripod gait and the m-tripod. Importantly, if flies were using the tetrapod gait, we would expect to see a cluster of data points around both the tetrapod at 0.6 and at 0.3. Both clusters are absent. Moreover, there is a tendency for the gait to become more pure tripodal with speed. In Figure 3a (top), GDIs were constructed on the basis of delays in stance start times; GDIs can also be constructed based on phase and also show that there is no change in gait (Figure 3a, bottom).

**Figure 3.**
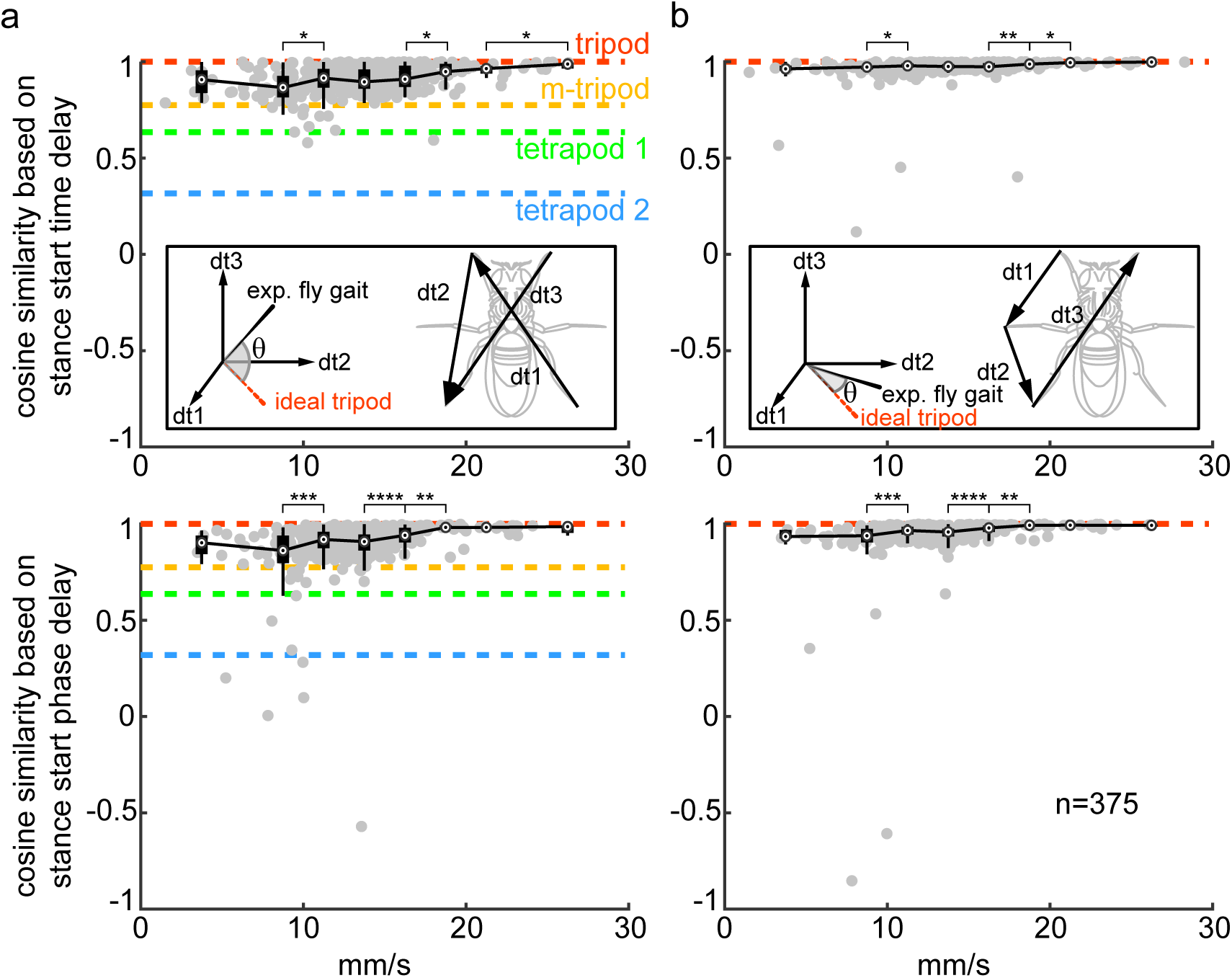
A new set of gait metrics, gait delay indices (GDIs), show that flies predominantly employ a tripod gait. (**a**) Top: cosine similarity of vector constructed from stance time delays [L1-R3, L3-R1, L3-L1] of experimental data to that of the same delays of an ideal tripod. Bottom: same as top but delays constructed from phase. Two sample Wilcoxin rank sum test shows a signficant increase in cosine similarities as speed increases.Only significant pairs are marked. (**b**) Top: cosine similarity of time deltadelays [L2-L1, L3-L2, L3-R1] of experimental data to ideal tripod. Bottom: cosine similarity of phase deltadelays of the same time deltadelays from the same sets of pairs.

We employed a second version of GDI which encapsulates how tripodal a gait is. Instead of considering how synchronized the legs within a tripod is – a measure that is likely to be erroneous because of small differences in phase between the legs in a tripod, we focus on the delays between legs that are part of opposing tripods: L1-L2, L2-L3, L3-R1 (Figure 3b). The majority (94%) of the data had a cosine similarity in the vicinity of one (Figure 3b). The results obtained from both delays and phases are similar, and once again do not support the idea that there is a speed-dependent transition from the tetrapod gait to a tripod gait.

In sum, GDIs encapsulate phase delays into a single metric which summarize gait. GDIs presented in Figure 3 show that a fly’s gait can be described as a modified tripod during which the prothoracic leg of the tripod is ahead of the other two legs in the tripod.

## Flies walk with a mid-stance maxima in CoM speed and height

Given that flies predominantly walk using a tripod gait, we analyzed flies’ CoM kinematics over a tripod gait cycle. Because the legs in a tripod are not synchronous, we defined the start of the tripod as the time point halfway between the very first foot landing, and the last foot of the preceding tripod lifting off; similarly, the tripod ends halfway between the last lift-off time of the tripod of interest, and the very first foot landing time of the following tripod (see dotted blue lines in Fig. 4a for definition).

**Figure 4.**
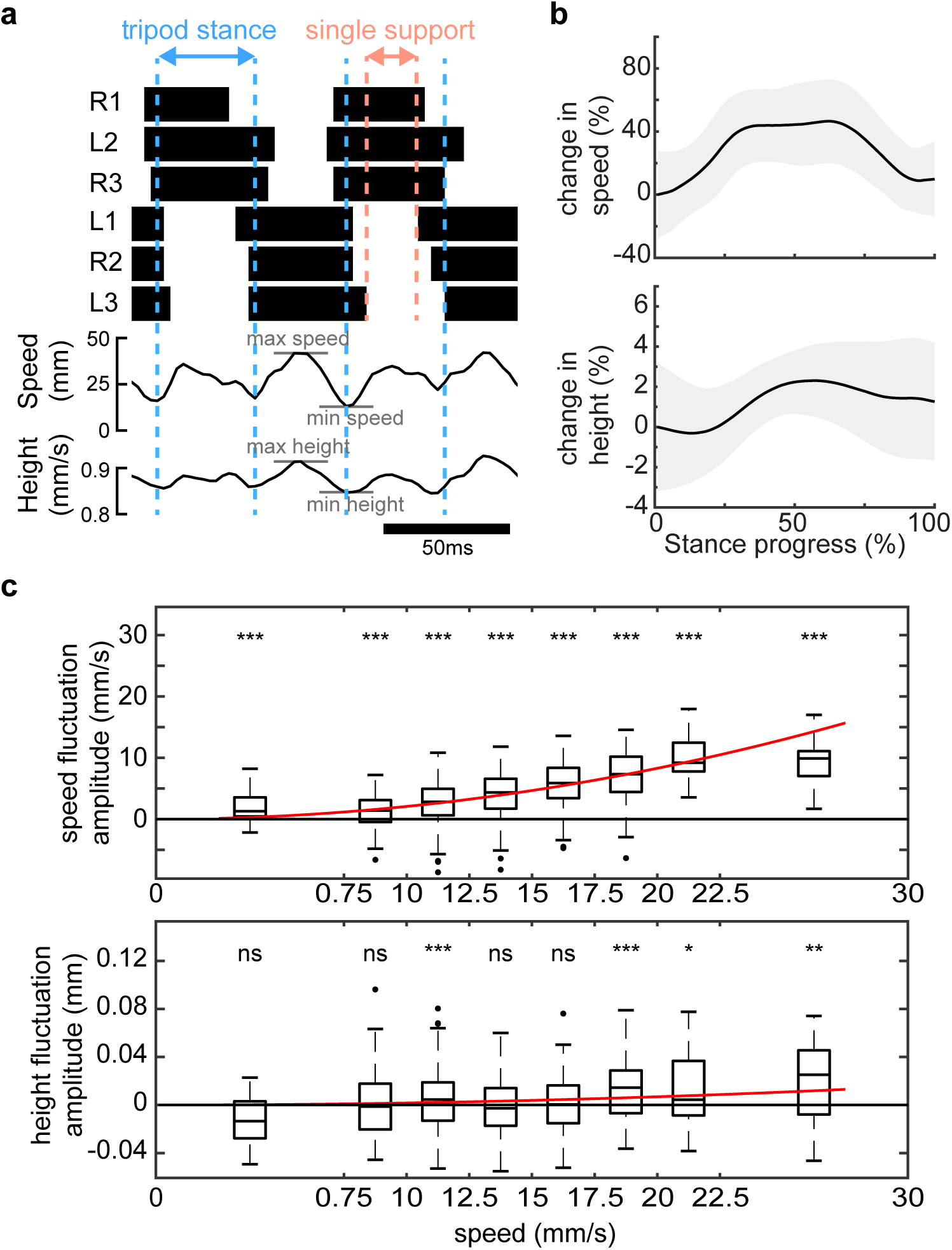
CoM kinematics during walking: CoM has a mid-stance maxima in height and speed. (**a**) Example trace showing the changes in CoM speed and height. Dotted blue lines are the boundaries between consecutive tripod stance (see text). Dotted orange lines are the boundaries of a single tripod stance. CoM has a clear mid-stance maxima in height and speed. (**b**) Top: average of normalized speed fluctuations during tripod stance phases. Shading: ±1 SD. Bottom: average trend of normalized height fluctuations. (**c**) Speed and height fluctuations increase with speed. Red lines are the best fit of y=ax^2^ to the data. Wilcoxin signed rank test to test whether there is a mid-stance maxima in speed and height.

As previously reported for stick insects^33^ and *Drosophila*^8^^,^^10^^,^ the horizontal speed of the CoM reached a maximum at mid-stance (Figure 4a). The mean trend is shown in Figure 4b (top); there is a 40% change in speed over each tripod. The change in speed is strongly dependent on the average speed (Fig. 4c top). The CoM height also has a mid-stance maxima (Figure 4a and 4b, bottom). The change is height is not as prominent or consistent as the change in speed.

It is important to note that a mid-stance maxima in speed is incompatible with most current mechanical models for locomotion which would yield a mid-stance minima in speed. Recently, we have proposed a model, angular spring modulated inverted pendulum (AS-IP), which can yield a mid-stance maxima in speed ^28^. One characteristics of this model is that the fly vaults over a stiff leg, and would therefore predict a maxima in the height of the COM, as we observe here. However, the fractional change in height is much smaller than that predicted using the AS-IP model. As will be elaborated in the next section, the smaller increases in CoM height compared to AS-IP suggests that the legs are compliant, not stiff.

## Angular and radial spring-loaded pendulum – a possible general mechanical model for insect walking

In a spring-loaded inverted pendulum (SLIP) model used to model the kinematics of a cockroach during running, it is assumed that all the mass of the cockroach is concentrated into a point mass which is supported by a single, massless, effective leg (Figure 5a). During the first half of stance, as the body moves through the stance phase, the spring is compressed, converting kinetic energy into elastic energy stored by muscles and tendons. During the second half of stance, the elastic energy stored in the leg spring is converted back into kinetic energy. Thus, the kinetic energy is at its lowest value at mid-stance, as is the height in most cases. This mid-stance minima in speed and height is also observed in the cockroach, making SLIP a simple, conceptual model for cockroach running. SLIP can also produce a mid-stance minima in speed at the same time as it produces a mid-stance maxima in height (Figure 5a). However, by its very nature, SLIP cannot produce the mid-stance maxima in speed observed in the fly (Figure 5a).

**Figure 5.**
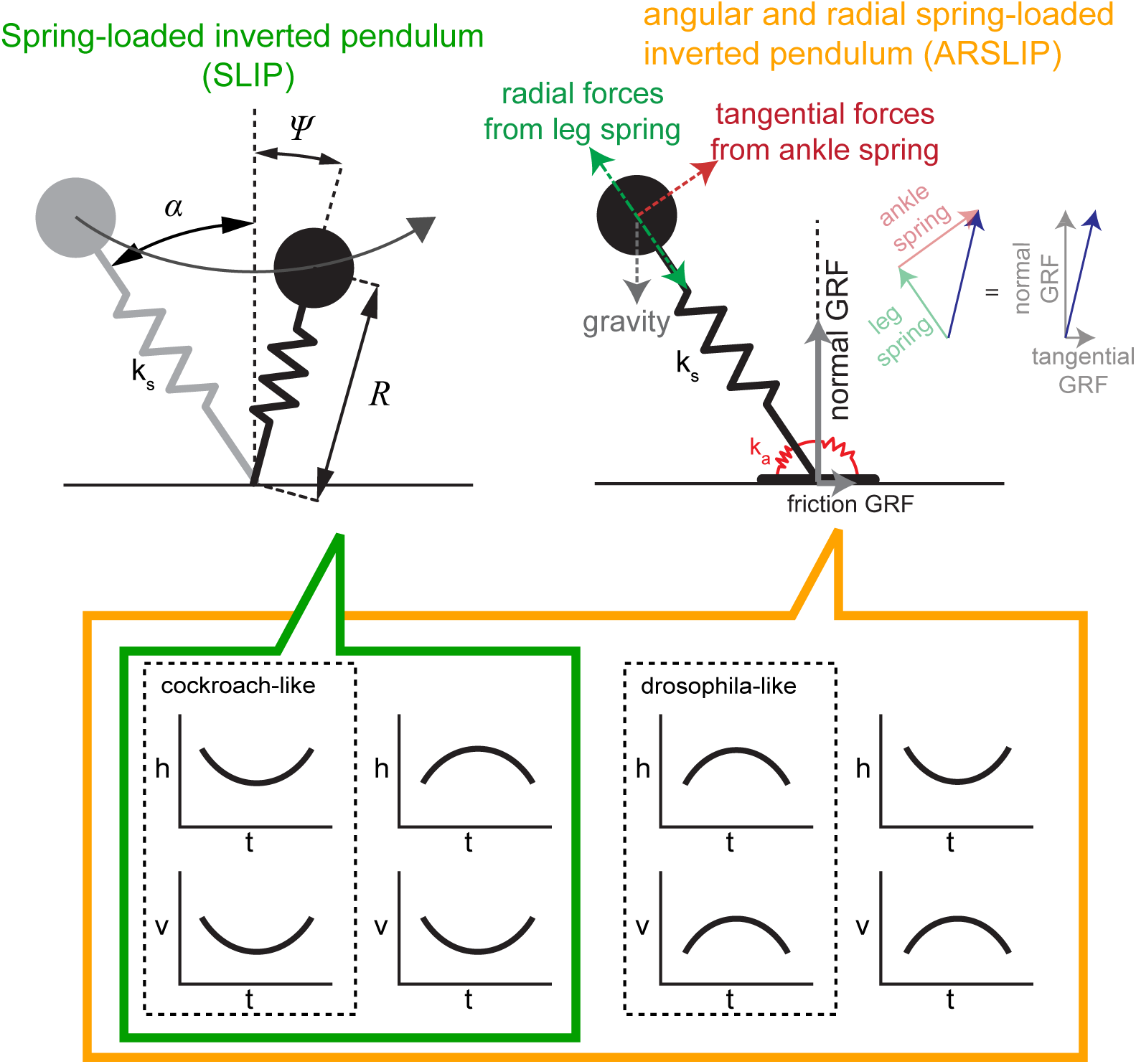
Limitations of SLIP and how they can be fixed by the proposed model - ARSLIP. Schematic showing SLIP - a simple model used to model legged locomotion in which all the stance legs are replaced by a single effective spring. SLIP can only produce two kinds of kinematic pattern. Kinematic pattern produced by fly cannot be modeled by SLIP. A simple modification of SLIP to add another spring that resists the legs movement from vertical results in the ARSLIP model. ARSLIP model can be employed to model fly’s locomotion. The angular spring is modeled as “ankle springs” for generality across animals including bipeds. The forces generated by the angular spring, radial spring, and gravity are shown with force vectors acting on CoM. Resultant ground reaction forces are shown as force vectors acting on the feet. SLIP will only produces forces marked with green arrows.

The fly-like mid-stance maxima in speed can be modeled by a simple extension to SLIP in which an angular spring resists the movement of the leg from the mid-stance position (Figure 5b). In Figure 9, we will discuss how insects can produce forces similar to the model proposed in Figure 5b simply by the virtue of three legs supporting the body at any moment. Moreover, insects also have a wide array of mechanism to generate attachment forces between the stance leg and the substrate^34^. These attachment forces are transmitted to the body as either forces along the leg or tangential to the leg. In the SLIP model, the ground reaction forces (GRFs) which ultimately move the body are always along the leg, and therefore forces tangential to the leg cannot be modeled. Adding an angular spring allows the transmittal of GRFs that are not along the leg – there is a component of the force along the leg, and a component tangential to the leg. Importantly, the angular forces reverse direction at mid-stance, as such they aid forward progression during the first half of stance, and oppose forward progression during the second half of the stance. This pattern is exactly opposite to the pattern created by SLIP. Depending on whether the leg spring dominates, or the angular spring, one can get a cockroach-like speed minima or fly-like speed maxima (Figure 5b). Thus, we hypothesize that each tripod can be replaced by an effective leg which functions as an **<u>a</u>**ngular and **<u>r</u>**adial **<u>s</u>**pring **<u>l</u>**oaded **<u>i</u>**nverted **<u>p</u>**endulum or ARSLIP.

## ARSLIP models the kinematics of a fly’s CoM during walking

To test the performance of the two models, SLIP and ARSLIP, we fit both the models to the CoM kinematics. Because there is considerable overlap between the stance times of the two tripods, a complete model would involve two effective legs, each of which functions as either SLIP or ARSLIP. A model with two effective legs would have too many parameters and might obfuscate many of clear insights that we obtain from modeling, therefore we modeled the CoM kinematics using a single effective leg. In contrast to most other work which employs simplified models as passive models, in this study these models are only passive at the timescale of a single step. In other words, we conceptualize the model parameters as an approximation of the control exerted by the fly at each step. In the ARSLIP model, at the beginning of each step, the fly chooses as its initial condition, angle of attack (*α*), angular speed (Ω), leg length (*R*), and radial speed (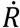). The evolution of the CoM depends on the mass (*m*), angular spring constant (*k_a_*), leg spring constant (*k_s_*), the natural leg length (*R_nat_*) and foothold location (*F*). The only difference in SLIP is that there is no angular spring, and hence *k_a_*.

We use an optimization algorithm to minimize the root mean squared error (RMSE) between the theoretical prediction of the position of the CoM predicted by SLIP and ARSLIP and the experimentally measured position. SLIP can model the small increase in the vertical position of the CoM which is a combination of two competing effects: there is an increase in CoM height due to progression of the CoM from its extrema to the vertical mid-stance position, and a decrease in the CoM height due to the compression of the leg spring (Figure 6a). But, as has been described previously^28^, SLIP fails to describe the horizontal evolution of the CoM. The failure of SLIP to describe the forward progression of the CoM is clear from the plot of the theoretical and experimental speed (Figure 6a, bottom panel): the theoretical speed profile has a mid-stance minima, whereas the experimental profile has a mid-stance maxima. In ARSLIP, the angular spring accelerates the CoM during the first half of stance phase, and decelerates the CoM during the second-half, and can therefore compensate for or even overcome the effects of SLIP. Therefore, the ARSLIP not only describes the vertical displacement of the CoM like SLIP, but also the horizontal displacement and the speed of the CoM (Figure 6b).

**Figure 6.**
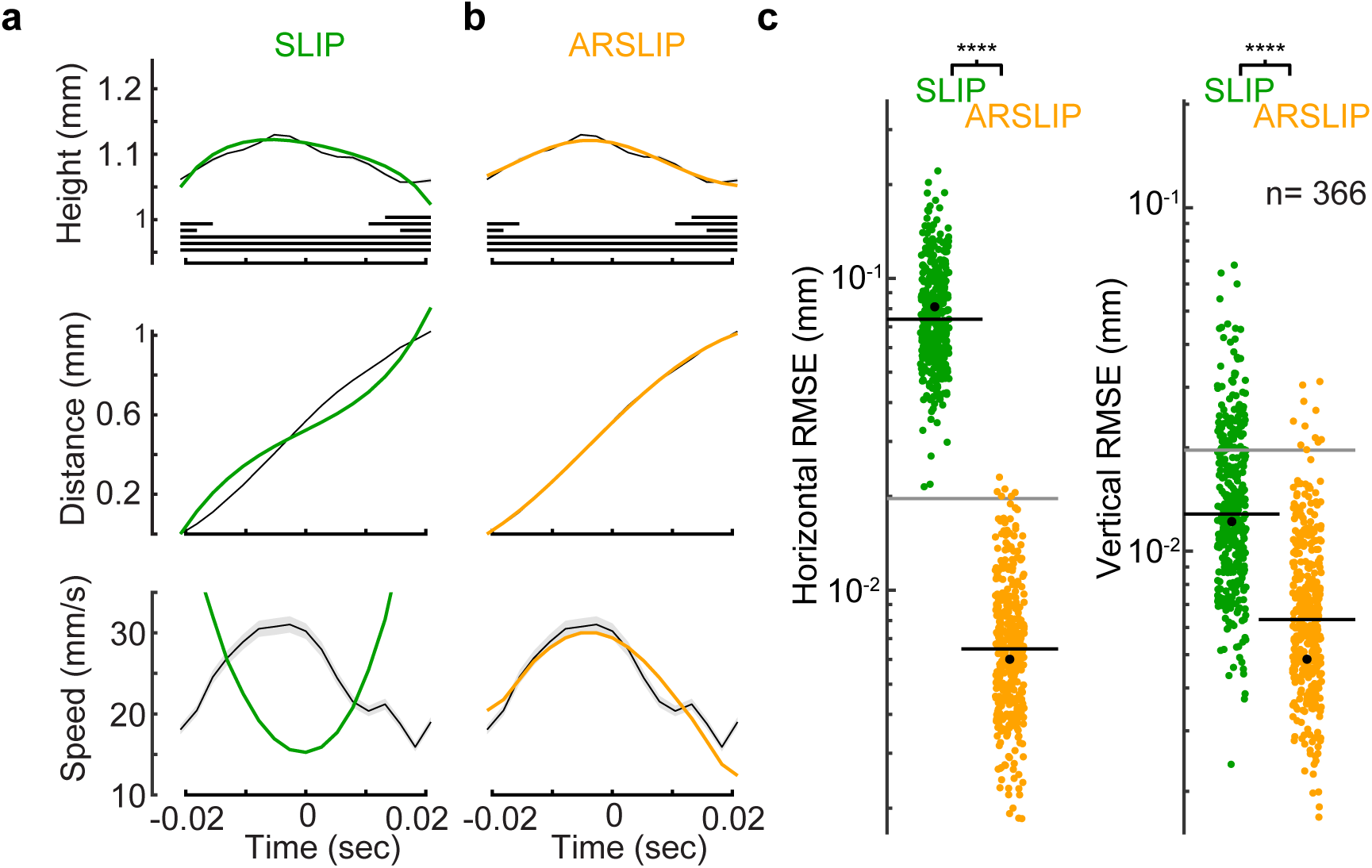
ARSLIP is an excellent model for describing the kinematics of a fly’s CoM. (**a**) Top panel: CoM height and the gait map. Middle: horizontal movement of the CoM. Bottom: speed of the CoM. The best fit of the SLIP model- to the height and horizontal position - is shown in green. Speed was not used to fit, but is plot to simply show why SLIP fails (**b**) Same as **a**, except that ARSLIP fits are shown. (**c**) ARSLIP is a significantly better model. The example step presented in **a** and **b** is marked as a black dot, and was chosen close to the SLIP median RMSE. Black horizontal line is the median. Gray line is a conservative estimate of the experimental error.

In all, we fit 456 steps. The RMSE for each step that we fit is shown in Figure 6c and shows that RMSE for both horizontal and vertical CoM displacement are significantly smaller for ARSLIP than for SLIP (using Wilcoxon signed-rank test; *p* < 0.001 for both horizontal and vertical movement).

## CoM kinematics are unaffected by inactivation of leg sensory neurons

Because ARSLIP describes the fly’s CoM kinematics, control of walking can be conceptualized as tuning the variables which define the evolution of the CoM under the ARSLIP model. This tuning can be through a combination of feedforward neural mechanisms, through sensory feedback or through mechanical properties. As a first step to understanding the mechanism, we sought to understand the effect of removing sensory feedback. To this end, we genetically silenced all leg sensory neurons ^8^ by blocking synaptic transmission through the expression of tetanus toxin (TNT) in these neurons. We will refer to the genetically silenced flies as *5-40^Leg^* flies (see methods and ^8^ for details).

Consistent with earlier findings, 5 − 40*^Leg^* flies used a tripod gait (Figure 7a). GDI also shows that the gait is close to a tripod (Figure 7b). The CoM kinematics – the CoM height and horizontal speed have the same trend as the wild type, and is at a maximum at mid-stance (Figure 7c). The increase in height and speed at mid-stance is more prominent for the mutant than it is for the wild-type (Figure 7c). The greater increase in height and speed is simply because the mutants walk at a higher speed than the wild-type (Figure 7e). When the change in height and speed is plotted as a function of speed, there is no difference between the mutant and control flies (Figure 7D). Given that the mutants have similar kinematics as the wild-type, it is not surprising that ARSLIP outperforms SLIP as a model for CoM kinematics (Figure 7F, using Wilcoxon signed-rank test; *p* < 0.001 for both horizontal and vertical movement).

**Figure 7.**
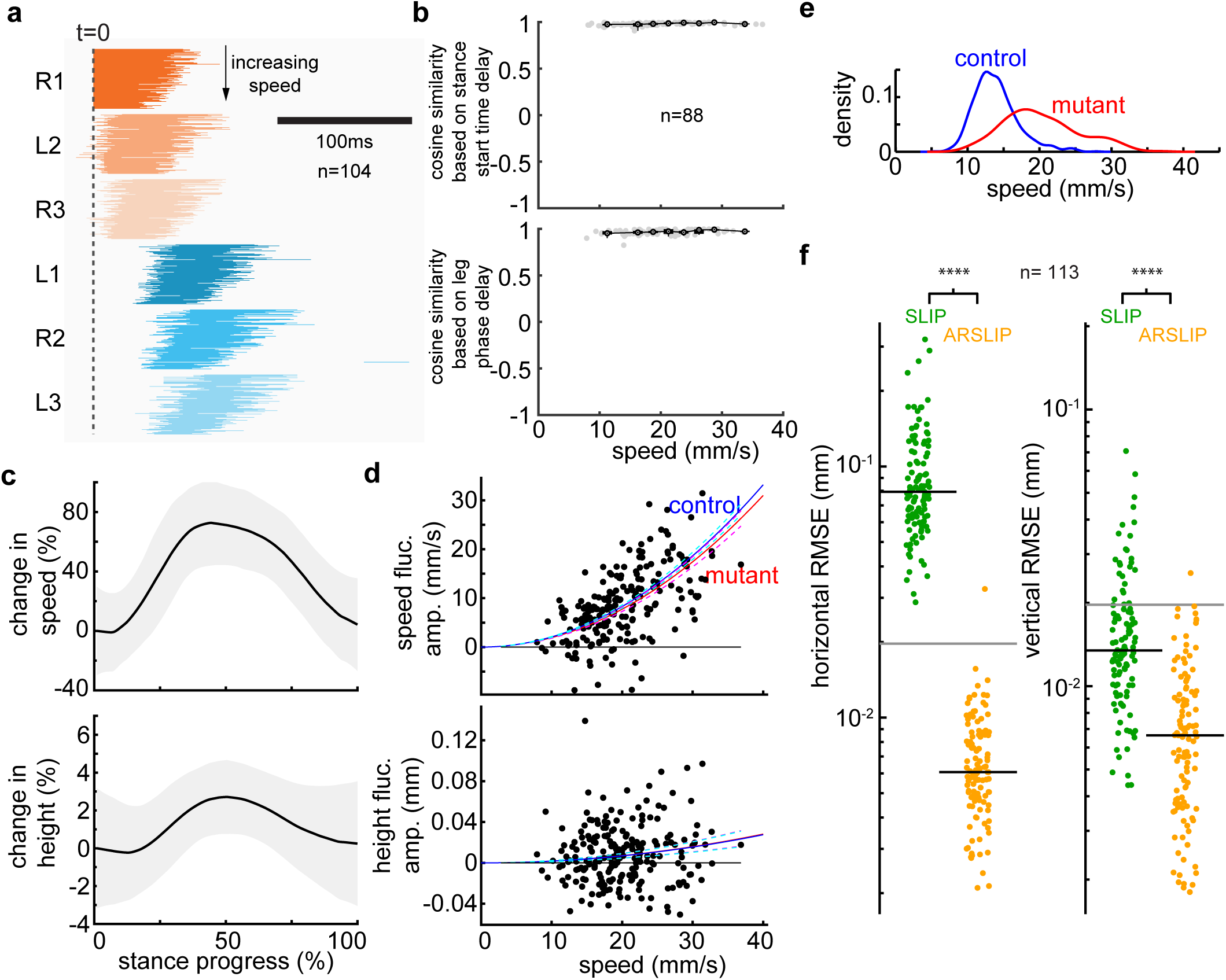
At a given walking speed, the kinematics of a mutant fly is virtually indistinguishable from control flies and is well described by the ARSLIP model. (**a**) Gait map plot analogous to Figure 2a shows that the gait is predominantly tripod. n =104. (**b**) cosine similarities (n = 88). based on delays ([L1-L2, L2-L3, L3-R1]) in stance start times (top) or phase (bottom). (**c**) Speed and height fluctuations during a tripod stance shows an increase in speed and height. (**d**) Height and speed fluctuation as a function of speed. Red line and blue lines are best fits of y=ax^2^ function to the mutants and controls respectively. The change in speed and height is similar for mutant and controls. (**e**) Mutants walk at higher speeds than controls. (f) ARSLIP is a signficantly better model than SLIP for describing a fly’s CoM trajectory (n=113).

That removing sensory feedback does not affect CoM kinematics is roughly consistent with the study which originally characterized the mutant^8^. The same previous study reported that the mutants walk slower than wild-type which, at first glance, is contradictory to our findings. However, the absolute speed with which the mutants walk is very similar in both studies. The difference in relative speeds between the mutant and controls is attributable to the fact that our control flies walk at speeds slower than the speeds reported previously. Because the mutants are strikingly similar to control flies in most respects, it is reasonable to treat the biomechanics of insect walking (straight walking on a smooth surface) as a combination of feedforward control of the limb, and the physics of the interaction between the stance legs and the external world. In the rest of this study, we will develop this idea further.

## An analysis of ARSLIP potential energy surface (PES) reveals a critical boundary that separates fly-like and cockroach-like gaits in the ARSLIP model

Although ARSLIP is a simple model, its parameter space can still support a large range of kinematic patterns; only a small region of this parameter space is employed in biological locomotion. To understand this parameter space, we will employ dimensionless variables:

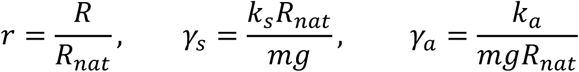

According to the ARSLIP model, forces at any time depends on *R (*or *r* in dimensionless terms), and Ψ (see Figure 5A). These forces cancel out at the fixed point of the system. It is easy to see that at mid-stance the forces tangential to the leg is 0: gravity acts along the leg and does not have any force directed tangential to it, and the angular spring does not exert any force. The forces perpendicular to the leg are gravity and the force due to the leg spring. These forces cancel out at a value of *R* where Eqn. 1 is satisfied.

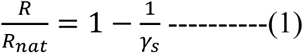

Therefore the fixed point of the ARSLIP is given by 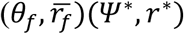 such that:

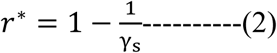

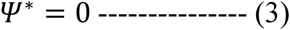

To assess how close to the fixed point a fly is at mid-stance, we plotted r as a function of *γ*_s_ (Figure 8A), and found, quite remarkably, that at mid-stance the flies CoM is right at the fixed point.

That the fly’s CoM passes through its fixed point at mid-stance considerably simplifies the analysis of the ARSLIP parameter space. One simplification is that its allows us to assess whether the CoM kinematics will be cockroach-like or fly-like by analyzing the PES around the fixed point (see Methods 3.6 for details). If the potential energy is locally minimum (or a stable equilibrium) at the fixed point, it follows from conservation of energy that the kinetic energy will be at a maximum at the same point, and we can infer a fly-like mid-stance maxima in speed. Conversely, a maxima in the potential energy implies a minima in speed and cockroach-like kinematics. Along the radial direction, a stable equilibrium node always exists, but in angular direction, the fixed point at *Ψ*\* = 0 can either be at a stable node or a saddle node. Therefore, whether the kinematics is fly-like or cockroach-like can be determined by evaluating the double derivative of potential energy along the *Ψ* direction. For a given mass, radial spring constant (*γ_s_*) and natural leg length, we can obtain *γ_a_* value (critical *γ_a_*) that satisfies

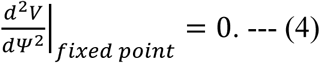

If the system has *γ_a_* greater than the critical *γ_a_*, the PES would have a local minimum at the fixed point (Figure 8b), and a mid-stance maxima in speed. If *γ_a_* is less than critical *γ_a_*, the PES would have a saddle point, and a speed minima at mid-stance (Figure 8b). The critical *γ_a_* boundary was obtained by combining Eqn. 1 with Eqn. 4.

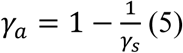

**Figure 8.**
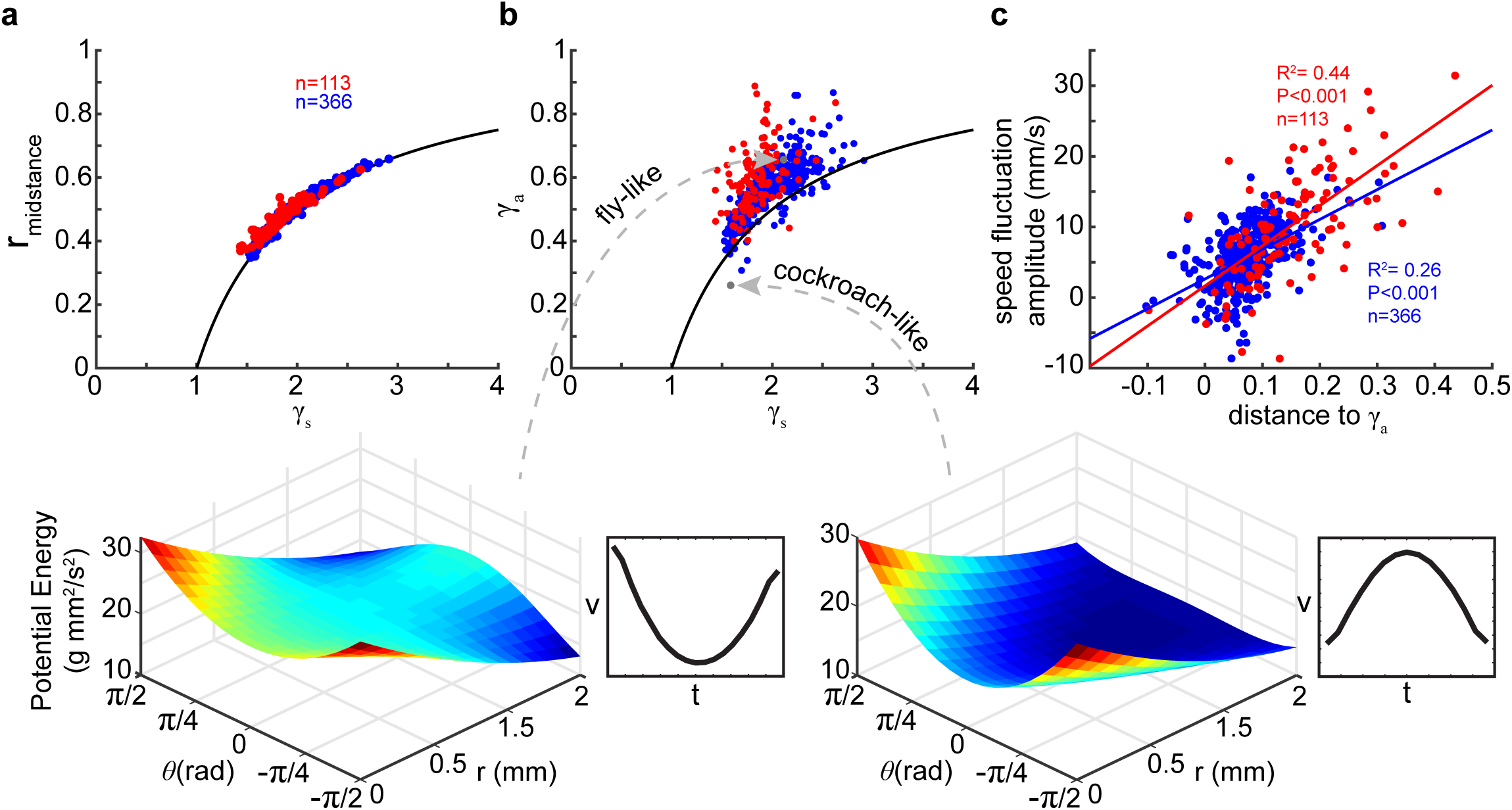
A critical boundary that separates fly-like kinematics from cockroach-like kinematics. (**a**) At mid-stance, flies CoM is close to the fixed point of the ARSLIP potential energy surface. (**b**) The line divides the ARSLIP surface into a region which will produce fly-like kinematics (above the line) and a region that corresponds to cockroach-like kinematics. The experimental fits lie just above the critical line (**c**) The deviation from the critical surface is predictive of the empirically obtained speed fluctuations.

With a few exceptions, all of our best fits of *γ_a_* were above the critical boundary consistent with the speed maxima in our dataset. Interestingly, most of the fits were close to the surface itself (Figure 8b). Is there a biomechanical reason? It turns out that the critical boundary has another relevance. The critical surface also represents (*γ_a_*,*γ*_s_) values at which 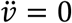: Due to conservation of energy in the system, Eqn. 4 implies Eqn. 6.

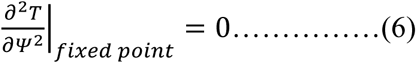

where *T* is a total kinetic energy. Since always increases with time, and the kinetic energy term is proportional to speed, Eqn. 6 implies 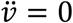. Therefore, for a given *γ_s_* value and a given angular amplitude, as one approaches the critical *γ_a_* the speed fluctuations become smaller, and low fluctuations in speed, in turn, allows the animal to walk at a relatively uniform pace. This argument is a generalization of the argument that was presented in an earlier work for a model which only had the angular spring and not the linear spring, a special case of ARSLIP with *γ_s_*→∞. For a given *γ_s_* value, the corresponding value of *γ_a_* at the surface will yield the least speed fluctuation for any arbitrary set of initial conditions. Consistent with this idea, the deviation in *γ_a_* from the critical *γ_a_* is predictive of the speed fluctuations during a step (Figure 8c).

## The simplest reduction of a point mass supported by three springy legs is the ARSLIP model

Thus far, we have shown that ARSLIP is an appropriate biomechanical model for insect locomotion. Can the insect legs produce forces that represent those produced by the ARSLIP model? To test this idea, we modeled the fly’s stance phase with the simplest biomechanical approximation of a tripod gait – a point mass representing all of the weight of the fly supported by three massless springy legs – which we will refer to as springy tripod (Figure 9a). In this model, the sagittal plane dynamics is governed by the sagittal plane projection of this tripod (red box in Figure 9a and details in Figure 9b). We compare how much of the kinematics of the step is represented by the geometry of the tripod of that same step (Figure 9c).

**Figure 9.**
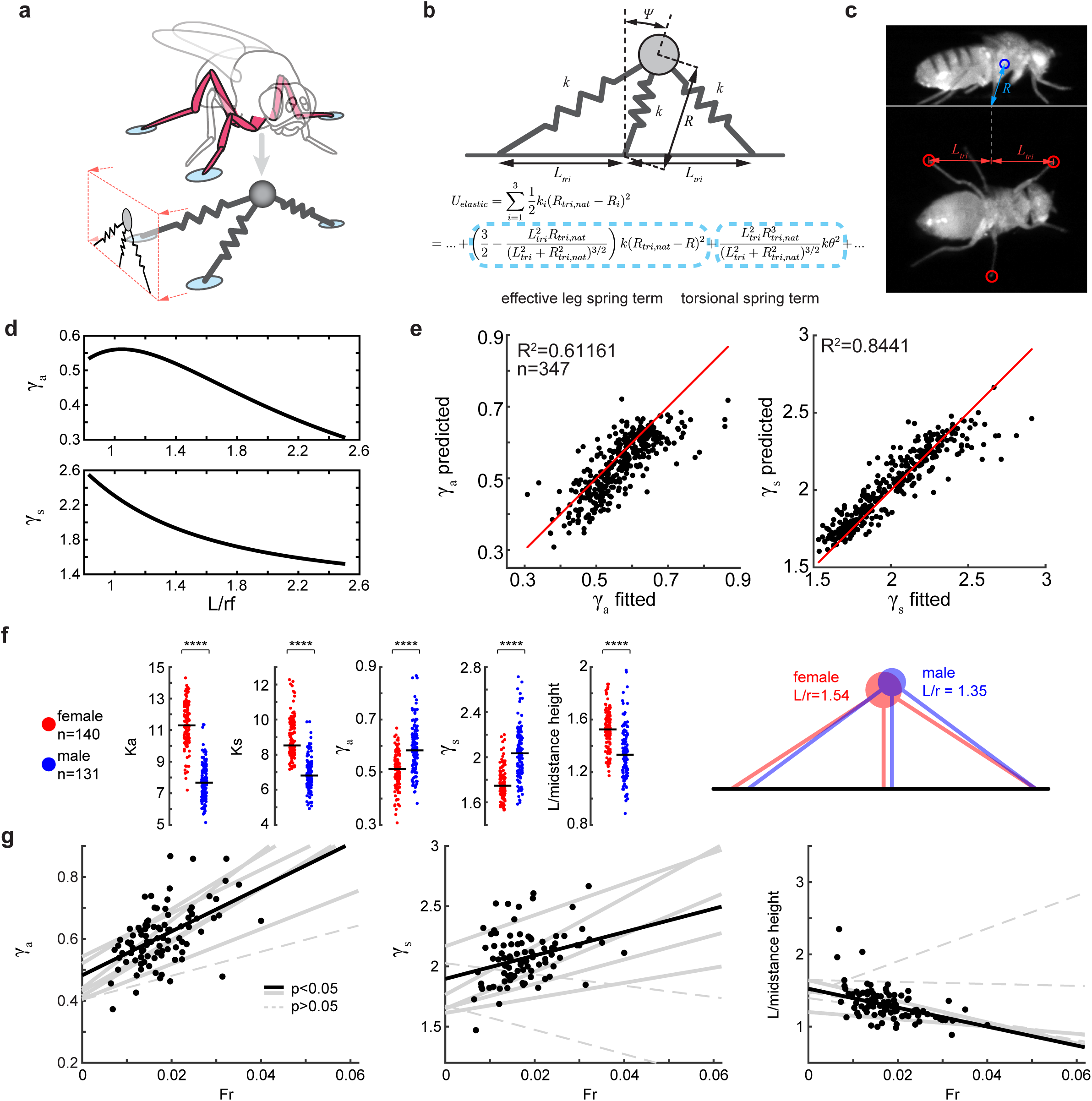
Actualization of ARSLIP using a springy tripod predicts the effect of tripod geometry on CoM kinematics. **(a)** Schematic of the springy tripod model. **(b)** Sagital plane projection of springy tripod reduces to the ARSLIP model. **(c)** We measure r_m_ and L for each step. **(d)** γ_a_ (top) and γ_s_ (bottom) decreases as L/r_m_ increases. **(e)** The geometry of the tripod correlates with the nondimensional spring constant (γ_a_: left and γ_s_: right). **(f)** Difference in tripod geomtry between males and females to account for their weight difference. Asterisks denotes p<0.00001 from Wilcoxon rank-sum test. **(g)** γ_a_ (left) and γ_s_ (middle) increase with speed in most flies. There is a corresponding decrease in L/r_m_. Solid lines are the regressions with p<0.05 from F-test, and dotted lines are p>0.05. Black dots (n=92) show values corresponding to individual steps for one fly.

First, to get insight into the potential energy produced by a system shown in Figure 9b, we considered the Taylor series expansion of the Lagrangian for this sagittal plane projection of the springy tripod (see methods for details). If, for simplicity, we assume that each leg has the same spring constant (*k*) and natural leg length (*R_tri,nat_*), then the potential energy of the system reduces to the equation :*C* + *A* · *k*(*R* − *R_nat_*)^2^ + *B* · *kΨ*^2^ (Figure 9b). The (*R* − *R_tri,nat_*)^2^ term corresponds to potential due to the leg spring term, and the *Ψ*^2^ term corresponds to the angular spring term. Together, these two terms correspond to the ARSLIP model, implying that the sagittal plane dynamics of a springy tripod is naturally described by the ARSLIP model.

Can a point mass supported by a springy tripod quantitatively produce the spring constants obtained from fits to the data? To test this idea, we used the empirically obtained tripod geometry (determined by *L_tri_* and *R_m_* in Figure 9c) and mass from our experimental data to determine the equivalent ARSLIP model that has approximately the same dynamics as the springy tripod around its mid-stance position (see Methods for details). Specifically, we obtained the following equations:

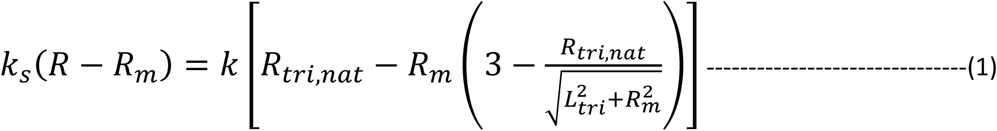

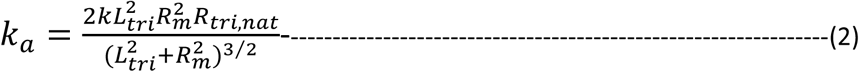

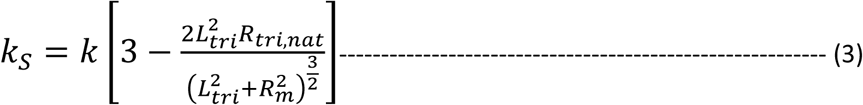

These equations relate the ARSLIP spring constants – *k_s_* and *k_a_* to *k*, the spring constant of each of the leg and the tripod geometry. The tripod geometry is defined by *R_tri,nat_*, the natural length of a fly’s leg and *R_m_*, *L_tri_* as defined in Figure 9b.

Everything else being the same, the spring tripod predicts that *γ_a_* and *γ_s_* will both vary over a two-fold range. As the ratio of *L_tri_*/*R_m_* increases both *γ_a_* and *γ_s_* decrease (Figure 9d). We can employ this dependence of *γ_a_* and *γ_s_* on the tripod geometry to examine how well the change in tripod geometry from one step to the next predicts the best fit *γ_a_* and *γ_s_* values from Figure 6. To this end, we found the best fit *k*(from Figure 9b); one constant *k* for each fly which best satisfies Eqns. (1-3). We found that the predicted *γ_a_* and *γ_s_* were close to the optimal *γ_a_* and *γ_s_* (Figure 9e). The springy tripod, despite all the simplifying assumptions, is able to describe the change in *γ_a_* and *γ_s_* with tripod geometry.

The analysis in Figure 9e shows that the spring constants that govern the CoM kinematics are strongly influenced by the tripod geometry. Does the fly change its tripod geometry systematically to control *γ_a_* and *γ_s_*? We examined two cases: First, we examined the differences between males and females (Figure 9f). Males are 50% lighter than females therefore the dimensional ks and ka values must be smaller. This is exactly what we observed (Figure 9f, leftmost two panels). The resulting non-dimensional *γ_a_* and *γ_s_* is actually larger for the male (Figure 9f) because males in our dataset walk faster (see Figure 9g). Thus, the males must be adjusting their *L_tri_*/*R_m_* ratio; this adjustment is exactly what we observe – males increase their *R_m_* decrease their *L_tri_* to obtain a smaller *L_tri_*/*R_m_* (Figure 9f, leftmost panel and schematic), and therefore walk with higher dimensionless spring constant.

Second, we examined how spring constants change with speed. We found that in a vast majority of flies, *γ_a_* and *γ_s_* increase with speed (Figure 9g). This increase in speed is usually reflected in a change in *L_tri_*/*R_m_*ratio (Figure 9g, rightmost column) implying that flies must be making their legs stiffer at higher speeds.

## Discussion

There are three main findings in this study:

- We describe a new metric for gait. This new metric shows that flies walk using a tripod gait for the entire range of speeds (Figure 2 and 3); the major change in gait as flies speed up is that the overlap between the two tripods decreases.
- The CoM speed and height increases to reach a mid-stance maxima. We show that a new model – ARSLIP – can not only model the kinematics of a fly’s locomotion, but has the flexibility to model locomotion that is characterized by kinematics dissimilar to flies.
- We show that the three legs of the tripod will naturally produce forces modeled by ARSLIP. The geometry of the tripod has a large influence on the forces exerted by the ground on the fly, and is modulated differently by males and females. The geometry of the tripod also contributes to change in speed.

These findings are discussed below.

Assessing gait is an essential first step for understanding locomotion. Recent progress in the acquisition of high-speed video, in automated detection of animal’s limbs, and in methods to assess inter-leg coordination has made it possible to amass data related to limb coordination from many insects^8,35,36^. However, because insects have six legs, each step is defined by five delays which describe the coordination of the five legs to a reference leg, and current methods employed to convert the five delays into a single gait metric are underdeveloped. GDIs employed in this manuscript represent a framework for distilling the five delays into a gait metric. We envision GDIs as a family of metrics which can be employed to assess different characteristics of an animal’s gait.

Based on GDIs, we conclude that flies predominantly employ a tripod gait during forward walking on a horizontal surface. Flies walking at low Fr numbers (Fr ~0.01), cockroaches running at high speeds (Fr >0.5), and ants walking at medium speeds (Fr ~0.3), all employ a tripod gait, implying that insects can employ the same gait across a large range of speeds. That insects employ a tripod gait across a range of speeds is in sharp contrast with the systematic transition from walk to trot to gallop in quadrupeds^37,38^. Gait transition in quadrupeds is well explained on the basis of energetics – for each speed there is an energetically optimal gait^38^. It appears that either the same relationship between gait and energetics do not hold in insects, or gait choice in insects result from constraints other than energetics. This interpretation is consistent with a recent study which suggests that the reason for the choice of tripodal gait in insects derives from a tripodal gait being a favored gait for climbing^39^.

Because a fly’s gait can be approximated by an alternating tripod, a tripod can be employed to ground both an analysis of the CoM kinematics, and biomechanical models of locomotion. In essence, as a first approximation, at any time a fly’s CoM is being propelled by the three legs of a tripod. In a previous study^40^ we have shown that walking slowly but uniformly, as flies do, requires tangential forces which reverse at mid-stance. In this study, we show that a springy tripod (Figure 9) will naturally produce such restorative forces, along with the normal spring forces that act along the leg. In the sense of “templates and anchors” idea^41^, ARSLIP is the template for fly locomotion, and this template is anchored by a springy tripod. An analysis of the ARSLIP parameter space provides two crucial insights: First, the height of the CoM at mid-stance is remarkably close to the fixed point (Figure 8A) of the ARSLIP system implying that at mid-stance there is no net force on the animal. Second, flies use a value of *γ_a_* such that the fluctuations in speed over a step is the smallest. Thus, the strategy a fly appears to employ is to choose a *γ*_s_ such that the fixed point at mid-stance, and *γ_a_* which ensures small velocity fluctuations during a step. Further work is required to understand the advantages of this strategy, and whether it represents an optimization either in terms of simplifying neural control, or in terms of stability. Conservatively, it appears to be a sensible strategy which will prevent large fluctuation in either height or speed.

An analysis of the springy tripod based on the measured geometry of the tripod shows that the step-to-step variation in the geometry of the tripod affects the CoM kinematics, and is an important mechanism by which the fly controls its trajectory. We present two scenarios in which the geometry of the tripod play a pivotal role: First, it appears to be an important mechanism by which males – who are ~50% lighter – achieve the same kinematics as females. Second, increasing the spring constants - *γ_a_* and *γ_s_*— by decreasing the spread of the tripod appears to be an important mechanism by which flies control speed within a step. These mechanisms of control of speed will not be observed in typical measures underlying control of speed such as step length or step frequency, and underscores the importance of mechanics in the control of speed.

It is possible that ARSLIP is a general model for locomotion. We have already shown in Figure 8 that different regimes of ARSLIP yield either fly-like or cockroach-like kinematics in terms of speed. As detailed in the companion manuscript (Antoniak et. al.), ARSLIP is also an excellent model for human walking. Human walking and fly walking reside in different regions of the ARSLIP feature space - humans walk with an extremely stiff leg spring compared to flies. Remarkably, even during human walking, the leg length is close to the fixed point at mid-stance (Figure 10 a). Moreover, because the *γ_a_* in the case of human walking is not proportionately larger, humans occupy a region of the ARSLIP that is below the critical surface (Figure 10 b) consistent with the predictions from our analysis in Figure 8. Thus, it seems possible that ARSLIP is a general model of locomotion. Further work is necessary to exhaustively test this hypothesis.

**Figure 10.**
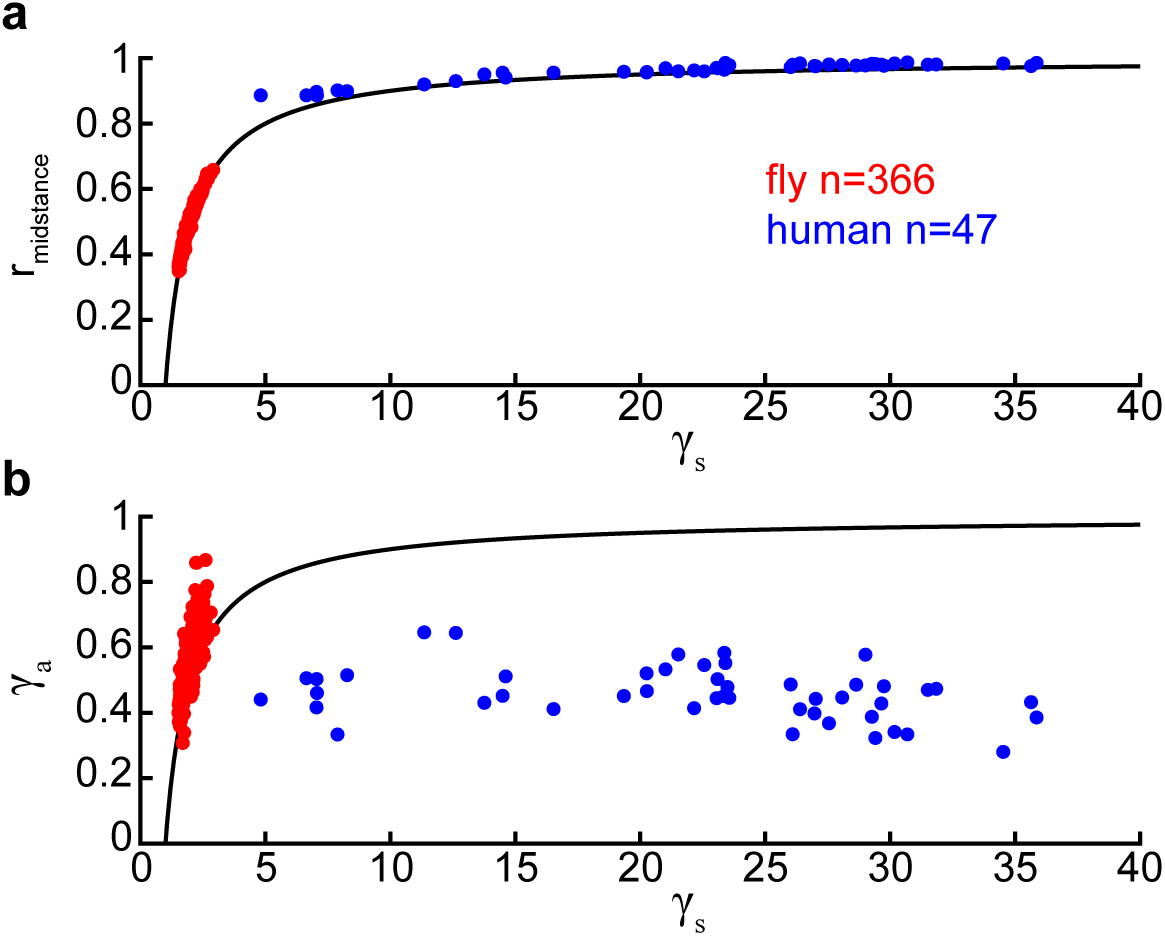
Best fits of ARSLIP to fly and human reveals underlying mechanical difference between the two speices. **(a)** Both fly and human have equilibrium leg length at midstance. Note that for a human, midstance leg length of ARSLIP was an initial condition, but for a fly, it needed to be solved. **(b)** Fly and human exist in different gamma regime.

## Methods

### 1. Flies

The flies were reared in an incubator that was maintained at 25°C, and 12 hr: 12 hr light:dark cycle. Ten minutes before experiment the flies were removed from a vial and their wings were removed using forceps on *CO*_2_ anesthesia.

Wild type strains were *w1118*, *Berlin K*, and *Oregon-R-C* (or Oregon C) (Bloomington stock numbers: 5905, 8522, and 5, respectively). We experimented with different wild-type strains because we wanted to record steps at a range of walking speeds, and working with multiple strains ensures that any general principle we discover is indeed general. Apart from the wild-type strain, we also experimented with flies whose synaptic transmission in the leg sensory neurons was genetically blocked. Flies of genotype 5-40-Gal4; dac-FLP was crossed with UAS-FRT-Stop-FRT-TNT were crossed in order to achieve this. In these flies, Gal4 was expressed in all sensory neurons but FLP recombinase was only expressed in the leg imaginal disc in order to activate tetanus toxin TNT, which blocks synaptic transmission.

### 2. Acquisition of behavioral data

Our experimental data consisted of the CoM position of the fly in all three dimensions, and the position of the fly’s footholds in the horizontal plane. In this section, we describe the procedure – both acquisition and processing – that yields this dataset.

#### 2.1. Recording chamber

The side walls and the ceiling of the chamber (inner *L* × *W* × *H*: 21×7×17mm) were built from microscope slides and ethyl-based instant adhesive (Loctite 495). A hole was drilled in one of the narrower side walls 1mm from the floor, to be used as an air jet nozzle for initiation of walking A coverslip of 0.13~0.17mm thickness was used as a floor of the chamber so that a distance between the side view and bottom view in a frame can be minimized. A thicker floor would have occupied unnecessarily large number of pixel rows, requiring a larger region of interest which would in turn decrease the frame rate. After a fly was placed inside the chamber, the chamber was secured on the coverslip using a tape. The chamber-coverslip assembly was then held horizontally using clamps. Below the assembly, a mirror was angled at 45 degree to the coverslip.. The mirror acted as a prism that redirected the bottom view of the chamber to the camera which was viewing from the side. The bottom and the side of the chamber were lit with infrared light, outside the visible spectrum of a fruit fly.

#### 2.2. Data acquisition

The procedure for data acquisition and processing were fully automated, except for a manual screening of raw videos before video processing. For a high frame rate and minimal optical distortion, USB 3.0 camera Basler acA1920-150um (380Hz at1024×779) and telecentric lens (Edmund Optics 0.40x SilverTL™) was used. This setup had MTF of 10% in vertical direction and 6% in horizontal direction at 25.39 lp/mm. he camera monitored the chamber at 30 Hz in real-time, until any motion within the field of view triggered acquisition at 380 Hz for 1.2 seconds. Exposure time per frame was set to 2.5 milliseconds. The motion was detected by measuring the difference between total pixel intensity values of the two most recent frames. Pulses of air jet were applied to the chamber to acquire more data and startle flies which would engage higher walking speed. For other trials, data was collected on spontaneous walks for normal or slower speeds. After each acquisition, the recorded video was saved to disk, if and only if the fly walked >5mm across the floor. This automated procedure could monitor and record a single fly for more than 10 hours.

#### 2.3. Tracking COM and the position of footholds during stance

The fly’s center of mass (CoM) was estimated by using the most prominent features of the fly as fiducial markers. The features were extracted on the first frame by using the minimum eigenvalue algorithm and following the extracted points throughout the video using Kanade–Lucas– Tomasi (KLT feature tracker in MATLAB ^25,42,43^. An estimated affine transformation matrix between the sets of feature points of consecutive frames was multiplied to the CoM position in the previous frame to estimate CoM position in the current frame (Supplementary Figure 3). Then, between every pair of consecutive frames, CoM was backtracked one step. The distance between original CoM and backtracked CoM is a reliable measurement of tracking error called forward-backward error ^44^. The distribution of errors is plotted in Figure 6. The error was small enough that we could evaluate the SLIP and ARSLIP models. The errors were also much smaller than the CoM trends (Supplementary Figure 4). The noise of the estimated CoM trajectories was insignificant, so numerical derivations of the trajectories returned velocity trends with a small noise (Supplementary Figure 4).

Location of footholds were automatically detected using a series of image processing algorithms detailed in Supplementary Figure 3. The basic idea was to binarize the bottom-view, and thin the resulting image to yield a skeleton. The end points of the resulting skeleton returned points including the actual footholds, along with other noisy or random points. The actual footholds were robustly detected by filtering out the noisy points, and extracting points that are located the furthest away from the CoM. The noise filtering was performed by removing small objects composed of less than 100 pixels. The legs were labeled based on the mean of each foothold trajectory in the CoM frame (details in Supplementary Figure 3).

### 3. Gait Analysis

We performed gait analysis using either the stance start times or instantaneous phase.

#### 3.1. Gait analyses based on a stance start times

For a gait quantification based on stance start times, a gait cycle was defined as a period during which all six legs had landed at least once. To identify all gait cycles in a given video, we first checked if tripod legs, i.e. [R1 L2 R3], were landing consecutively without particular order. If the tripod legs were landing consecutively and the other tripod legs, i.e. [L1 R2 L3], were immediately following without particular order, then we identified the cycle as a tripod gait cycle. For a set of leg landing sequence that could be defined as a cycle but could not as a tripod gait cycle were identified as unregistered gait cycle. The identified cycles were used for composite gait map analysis (Figure 2a, Figure 7a) and gait quantification analyses (Figure 3a top, Figure 3b top, Figure 7b top). For quantifying a gait based on stance start times, we compared stance start times of an experimental and synthetic tripod gait cycles. A synthetic tripod gait cycles had a same duty factor as an average duty factor of experimental gait, but with legs within tripod completely in-phase and the legs between tripod completely out of phase by half a cycle. For our gait metric, we first defined stance start time difference between a pair of legs as delta. A set of three deltas, L1-R3, L3-R1, and L3-L1, were chosen for distinguishing between tripod and tetrapod gait. Another three deltas, L1-L2, L2-L3, and L3-R1, were chosen for quantifying gait based on delays between legs that are out of phase in a tripod gait. Each set of deltas were measured for both experimental data and synthetic tripod and then these two deltas from experimental and synthetic gait were represented as three-dimensional vectors. Finally, a cosine similarity between the vectors was our gait metric. Cosine similarity is essentially a cosine of an angle between two vectors.

#### 3.2 Gait analysis based on leg phases

In a body coordinate system where origin is located at the CoM and the poisitve-y axis points to the anterior of the body, *y_i_*(*t*) of leg positions were measured. Since we only know leg positions during stance, we performed a linear interpolation of *y_i_*(*t*) during swing. To get instantaneous phase angles of the legs, Hilbert transform was performed on the times series data ^29,32,45,46^. Hilbert transform takes real-valued signal (a signal of interest) and returns complex-valued analytic signal which is used for accurately computing instantaneous magnitude and frequency of the real-valued signal ^47-49^. The time dependent angle of complex number of analytical signal is instantaneous phase angle.

For phase shift analysis shown in Figure 2d, an instantaneous phase shift relative to R1 was averaged over a single gait cycle. Here, a gait cycle was defined as a period when R1 completes a single full cycle starting and ending at a lift off phase angle (∅ = −*π*). For gait quantification shown in Figure 2a bottom, Figure 2b bottom, and Figure 6b bottom, phase angles of a synthetic tripod gait were generated for each experimental gait cycle. First, we generated *y_i_*(*t*) of the synthetic tripod gait to get its phase angles. To determine duty factor of the synthetic tripod gait, we took values from the regression for stance and swing durations at an average speed of the experimental gait cycle (Figure 1d). Travel distance of the stance phase leg of the synthetic tripod at a given speed was also determined from a regression fit of experimental data. We assumed that the legs within a tripod had the same *y_i_*(*t*), whereas the legs in the other tripod had *y_i_*(*t*) shifted by half a cycle. Hilbert transform of these time series returned the phase angles of the synthetic tripod. The method for getting cosine similarity between experimental gait and synthetic gait based on leg phases was the same as the one based on stance start time, except that instantaneous phase shift between a pair of legs was delta for this analysis. We took average of the instantaneous cosine similarity over a single cycle to get the final gait metric based on leg phases.

#### 3.3 Calculation for change in height and velocity

Height and speed change during stance phase was calculated as described below. s A time series of height or speed over a tripod stance was detrended by a line that connected the values at the beginning and end of the stance phase. Finally the maximum and minimum values of the detrended data were summed to get the change in height or speed value (Supplementary Figure 5).

#### 3.4 System of ordinary differential equations (ODEs) for SLIP and ARSLIP and details regarding fitting ODEs to individual steps

The following system of ODEs (Eqn. 4 and 5) of ARSLIP was derived using Euler–Lagrange equations. Polar coordinate system was chosen for simplicity in expression.

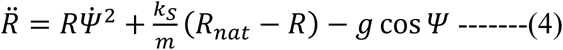

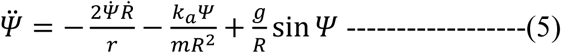

where *R*(t) is a leg length, *Ψ*(*t*) a leg angle from vertical axis, *k_s_* a leg spring constant, *k_a_* an angular spring constant, *m* a total mass of a fly, and *R_nat_* a natural leg length. Dot denotes time derivative. Detailed derivation of the ODE is presented in Supplementary section of Antoniak et al 2018.

For finding the best fit of ARSLIP trajectory to a given experimental trajectory, Global Search algorithm from MATLAB Global Optimization Toolbox was used ^50^. Objective function subject to minimization was RMSEs between time series trends for height and distance was summed and the summed value was used an objective subjected to minimization.

*K_s_* and *k_a_* were searched with the following inequality constraints (Eqn. 6 and 7):

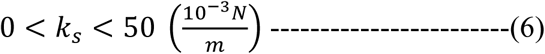

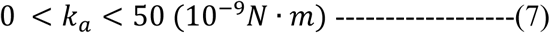

These bounds were large enough to that none of the fitted parameter values had the boundary value. *R_nat_* and *m* were experimentally measured from the strains (Table 1). Multiple middle leg length measurements of a fly were averaged (*R_real_*_,*nat*_), and then based on the average value optimal *R_nat_* was estimated by searching within ±10% boundary of *R_real_*_,*nat*_. *m* was measured by averaging weights of 10 flies of same species and gender on a scale (Mettler Toledo XP26; 0.001mg readability).

**Table 1.** Measured parameter values for each species and gender.

| fly<br>number | gender | strain | $R_{real,nat}$<br>(mm) | mass<br>(mg) |
| --- | --- | --- | --- | --- |
| 1 | female | Berlin K | 2.1032 | 1.0809 |
| 2 | female | Berlin K | 2.184 |  |
| 3 | female | Berlin K | 2.1031 |  |
| 4 | female | Berlin K | 1.9988 |  |
| 5 | female | Berlin K | 1.9869 |  |
| 6 | male | Berlin K | 1.8888 | 0.6798 |
| 7 | male | Berlin K | 1.9272 |  |
| 8 | male | Berlin K | 2.0646 |  |
| 9 | male | Berlin K | 2.0404 |  |
| 10 | female | W1118 | 2.042 | 1.123 |
| 11 | female | Mutant | 2.1359 | 0.9687 |
| 12 | female | Mutant | 2.1136 |  |
| 13 | female | Mutant | 1.8913 |  |
| 14 | female | Mutant | 1.9357 |  |
| 15 | female | Mutant | 2.1105 |  |
| 16 | male | Mutant | 2.0237 | 0.6224 |

Because a model with two legs would have too many parameters and would obfuscate many of clear insights that we obtain from modeling, we chose steps for which at least 25% of the time was spent with only the legs of a single tripod on the ground (see Methods for details). This criteria does not mean 75% of the step is spent with both tripods on the ground. Because the legs of the tripod are not synchronized, much of the time that is spent with both tripods on the substrate is the time it takes for a subset of legs from the other tripod to leave the ground. Initial conditions of 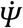 and 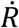, and were also measured from experimental data but optimal initial condition was searched within ±10% boundary of the measurements. Since we set the foothold location of ARSLIP as middle of front and hind leg foothold positions, initial conditions of *R* and *Ψ* could be determined from experimental data.

The same method was applied for fitting SLIP model, with the same conditions for the parameters except for *K_a_* due to lack of angular spring. The system of ODEs for SLIP is presented below.

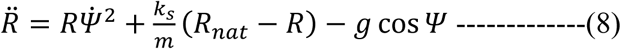

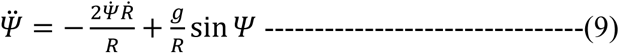

For both models, *g* value of 9.807*m*/*s*^2^ was used.

#### 3.5 Derivation of the critical boundary

In order to have maximum velocity at mid-stance (*Ψ* = 0), CoM should experience positive acceleration before mid-stance and negative acceleration after mid-stance. Parameter space that allows this behavior could be found using formula for non-dimensional horizontal acceleration (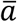) of ARSLIP (Eqn. 10).

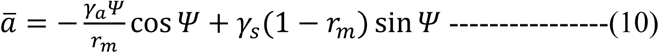

Where *r_m_* was a dimensionless leg length at mid-stance. The acceleration term will have opposite signs before and after the mid-stance with zero at the mid-stance. In the case for maximum velocity at mid-stance, we applied small angle approximation and constrained 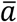 to be negative for positive *θ*. This resulted in a parameter space defined by *γ_a_*, *γ_s_*, and *r_m_* (Eqn. 11-13).

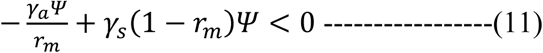

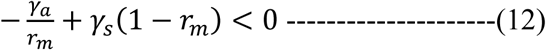

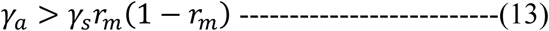

Therefore the following was a critical boundary that separated the two velocity trends (Eqn. 14).

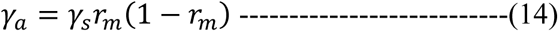

However, *r_m_* required solving ODEs, so the formula could not be used for predicting velocity trend, unless mid-stance was initial condition. Therefore, 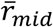 was approximated to fixed point value of r at mid-stance (*r*\*). *r*\*is a resting dimensionless spring length when the spring is standing vertically (mid-stance or *Ψ* = 0) against gravitational force. The following derivation of *r*\* begins with dimensional terms (Eqn. 15-17).

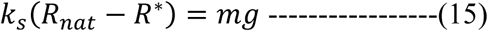

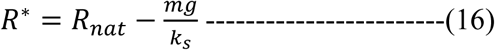

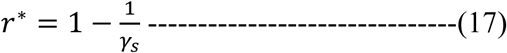

Substituting *r_m_* with *r*\* in the critical boundary formula (Eqn. 14), we get the following formula (Eqn. 18).

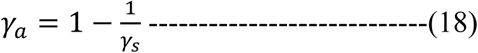

Given *γ_s_*, if a model’s *γ_a_* is greater than *γ_a_* from this formula, the model will experience maximum velocity as long as initial condition satisfies the constraint that *r_m_* will be close to*r*\*. In case of the fits for drosophila, *r_m_*tends to be close to match *r*\* (Fig. 2 methods).

#### 3.6 Derivation of formula relating tripod model to ARSLIP

The tripod model’s total elastic potential is given by

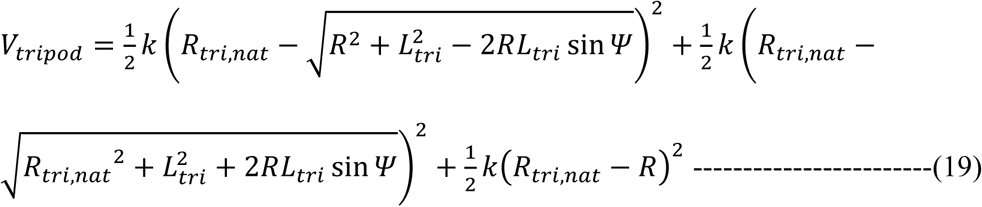

We claim that *V_tripod_* is equivalent to the ARSLIP potential for evolution that are close to the mid-point, *R* = *R_m_* and *Ψ* = 0.

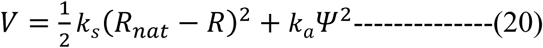

Since |Δ*R*/*R*_*m*|~0.1 and |*Ψ*|~0.2, it should be sufficient to show that the two potentials agree with each other up to quadratic order in fluctuations around the fixed point. This means that we need to ensure that the first and second derivatives with respect to *R* and *Ψ* at *R* = *R_m_* and *Ψ* = 0 are the same for both the potentials. Specifically,

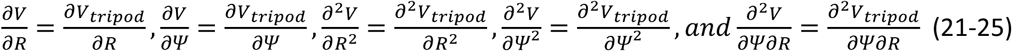

are the same for both the potentials. We note that the relations involving the first derivative of *Ψ* (Eqn. 22), and cross double derivative involving both *Ψ* and *R* are automatically satisfied (Eqn. 25). This shows that our assumption about the independence of the radial and angular springy forces are actually satisfied in the simplest tripod model.

Next to investigate whether the springy tripod model actually predicts the dependence of spring constants on tripod geometry, we are left with three equations, involving the first derivative of *r* (Eqn. 21), and the two double derivatives w.r.t. *R* and *Ψ* (Eqn. 23, 24). We have three parameters in the effective ARSLIP potential, *R_nat_, k_s_, and k_a_*, and thus it should be possible to satisfy all these conditions. Specifically we obtain the following equations:

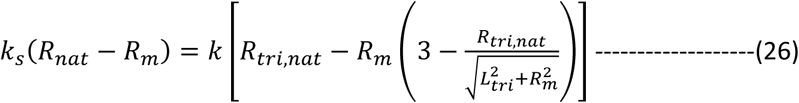

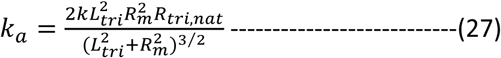

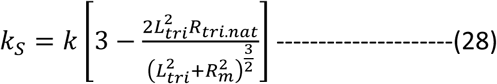

Thus we see that all the parameters of the effective potential, *k_s_, k_a_* and *R_nat_* can be determined in terms of the tripod potential parameters, *k* and *R_tri,nat_* and the geometric quantities *R_m_*, *L_tri_* (Eqn. 26,27,28). We obtained *R_m_* and *L_tri_* from the geometric data for each step. We assumed that a given fly has the same natural length *R_tri,nat_* and *k* and try to fit *k_a_*, and *k_s_* for all the steps of the given fly. This way we will obtain a best fit value of *k* and *R_tri,nat_*, but most importantly we can see whether the theoretical fitting provides the general experimental trend.

## Acknowledgments

We would like to acknowledge the members of Bhandawat lab for critical comments on earlier versions of the manuscript. This research was supported by NIDCD (VB), NINDS (VB) and an NSF CAREER award (VB).

## Author contributions

VB: Conceptualization; Supervision; Funding acquisition; Writing; analysis. CC: Conceptualization; Experimentation and analysis; Writing. TB: Conceptualization; analysis; Writing;

---

**Table 1:**
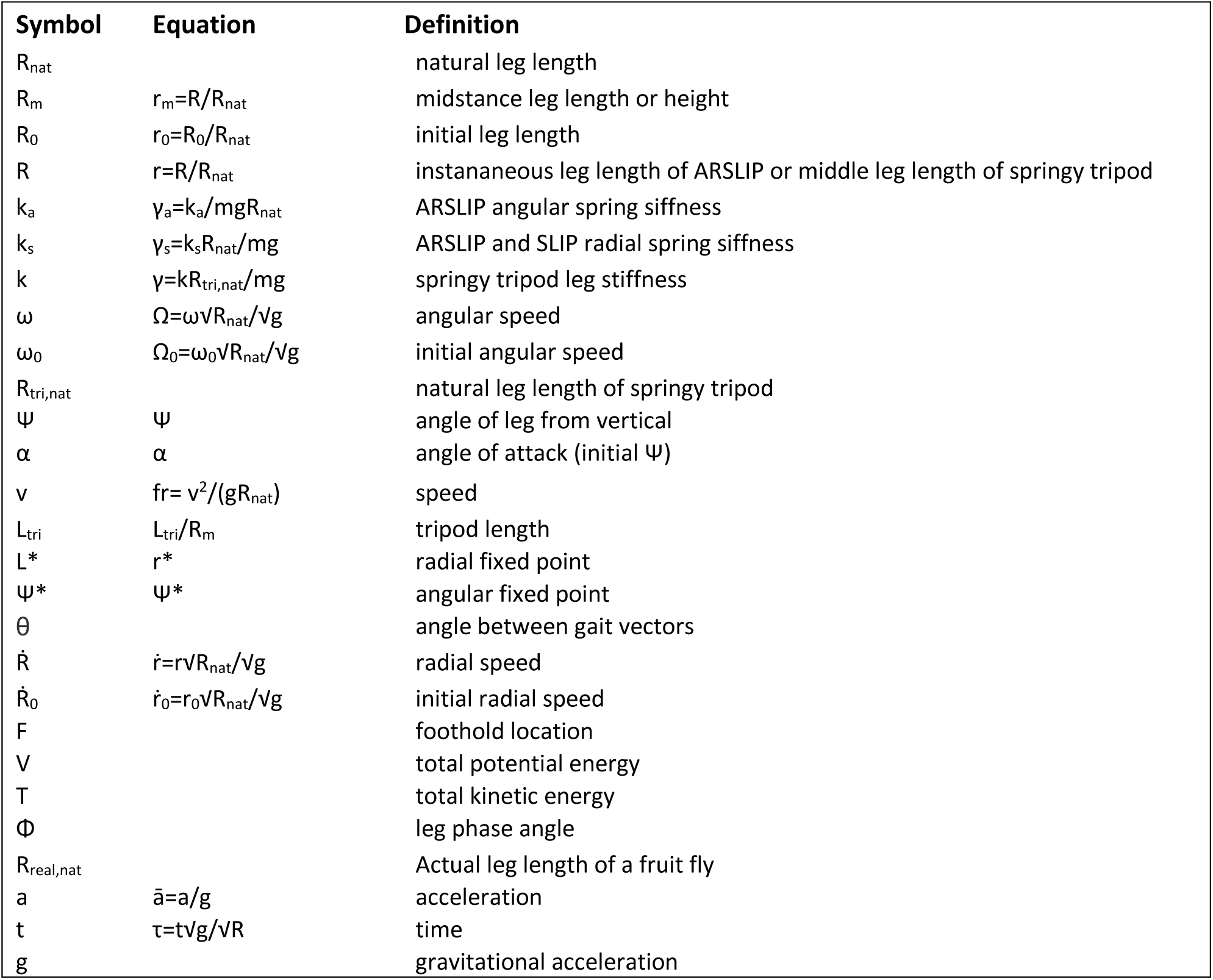
List of symbols.

| Symbol | Equation | Definition |
| --- | --- | --- |
| $R_{nat}$ | | natural leg length |
| $R_m$ | $r_m=R/R_{nat}$ | midstance leg length or height |
| $R_0$ | $r_0=R_0/R_{nat}$ | initial leg length |
| $R$ | $r=R/R_{nat}$ | instantaneous leg length of ARSLIP or middle leg length of springy tripod |
| $k_a$ | $\gamma_a=k_a/mgR_{nat}$ | ARSLIP angular spring siffness |
| $k_s$ | $\gamma_s=k_sR_{nat}/mg$ | ARSLIP and SLIP radial spring siffness |
| $k$ | $\gamma=kR_{tri,nat}/mg$ | springy tripod leg stiffness |
| $\omega$ | $\Omega=\omega\sqrt{R_{nat}/g}$ | angular speed |
| $\omega_0$ | $\Omega_0=\omega_0\sqrt{R_{nat}/g}$ | initial angular speed |
| $R_{tri,nat}$ | | natural leg length of springy tripod |
| $\psi$ | $\psi$ | angle of leg from vertical |
| $\alpha$ | $\alpha$ | angle of attack (initial $\psi$) |
| $v$ | $fr= v^2/(gR_{nat})$ | speed |
| $L_{tri}$ | $L_{tri}/R_m$ | tripod length |
| $L^*$ | $r^*$ | radial fixed point |
| $\psi^*$ | $\psi^*$ | angular fixed point |
| $\theta$ | | angle between gait vectors |
| $\dot{R}$ | $\dot{r}=r\sqrt{R_{nat}/g}$ | radial speed |
| $\dot{R}_0$ | $\dot{r}_0=r_0\sqrt{R_{nat}/g}$ | initial radial speed |
| $F$ | | foothold location |
| $V$ | | total potential energy |
| $T$ | | total kinetic energy |
| $\Phi$ | | leg phase angle |
| $R_{real,nat}$ | | Actual leg length of a fruit fly |
| $a$ | $\bar{a}=a/g$ | acceleration |
| $t$ | $\tau=t\sqrt{g/R}$ | time |
| $g$ | | gravitational acceleration |

**Supplementary Figure 1.**
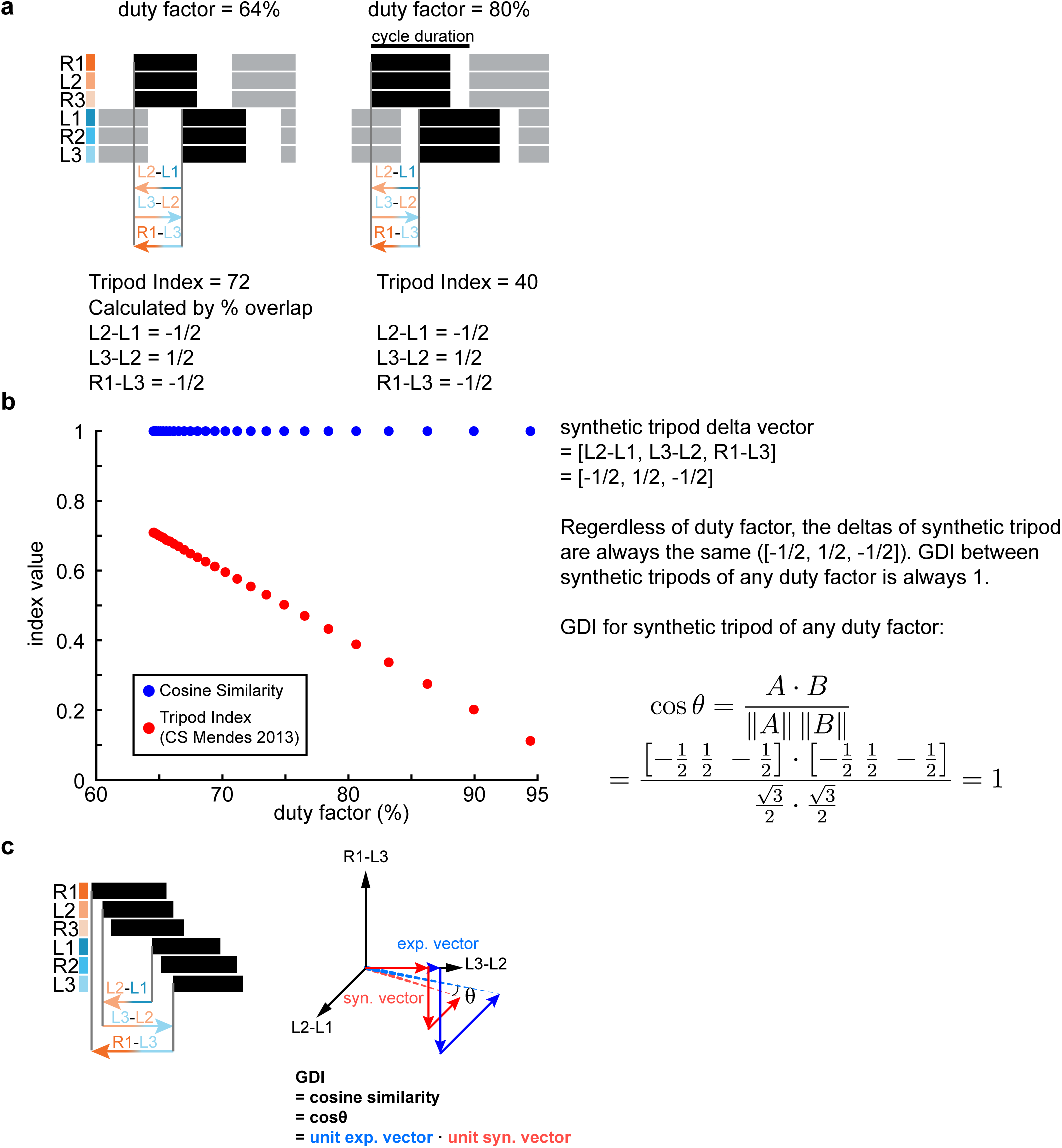
Gait metric comparison. **(a)** Synthetic tripod gaits with 64% duty factor and 80% duty factor. Although the absolute phases are a function of duty factor, the relative phases are not. One way to capture this idea is to create a unit vector using these delays. This unit vector is always [-1, 1, -1]/3^1/2^ for a tripod gait. **(b)** Effect of duty factor on our gait metric (blue dots) and tripod index by Mendes and his colleagues (red dots) (Mendes et. al. 2013). **(c)** Visualization of calculation of GDI.

**Supplementary Figure 2.**
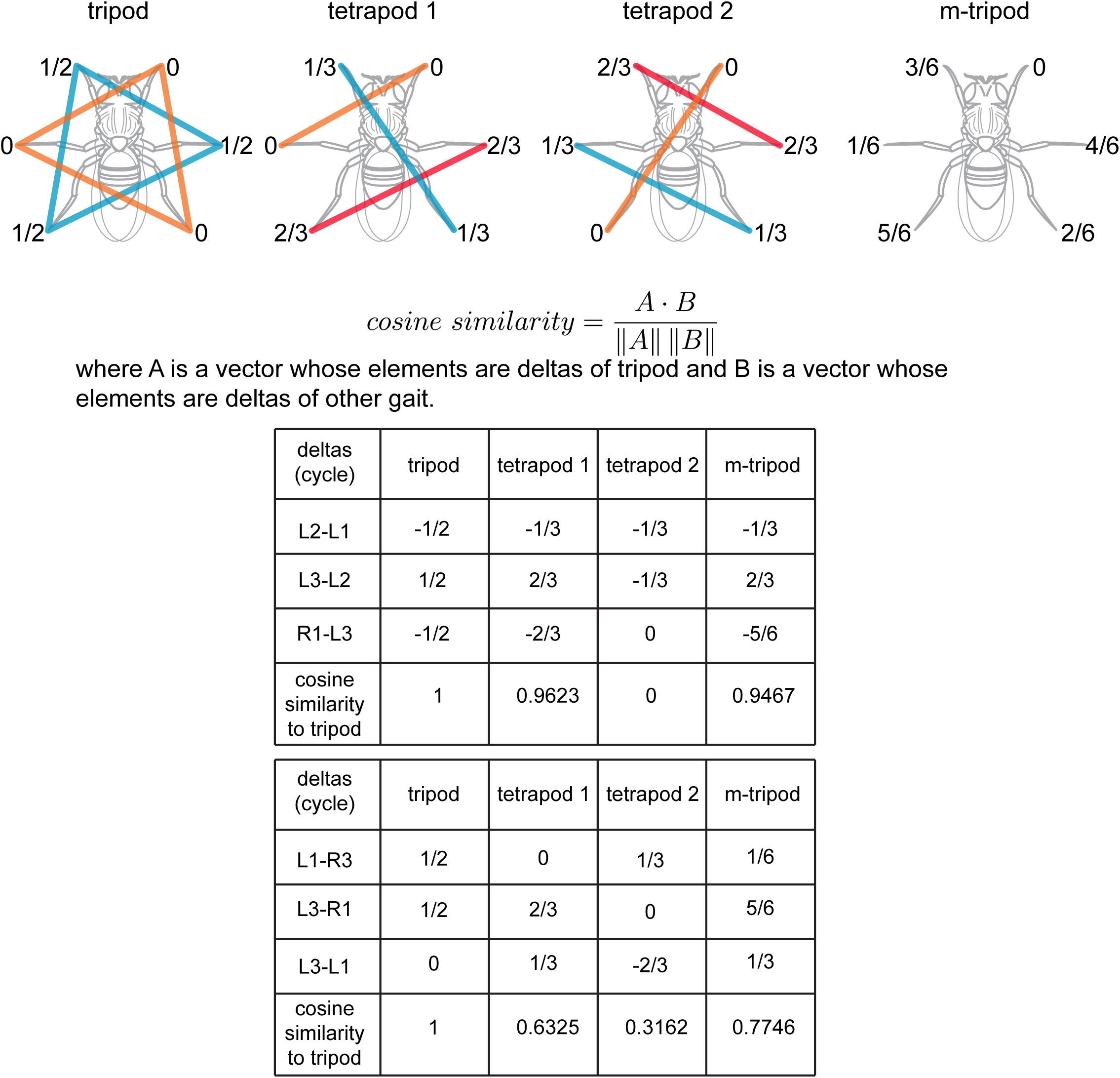
Phases/time delay between legs during three different gaits. Relative phases of the legs during tripod, tetrapod 1, tetrapod 2, and m-tripod as conventionally defined. Here, phase value ranges from 0 to 1. 0~1 is a complete cycle.

**Supplementary Figure 3.**
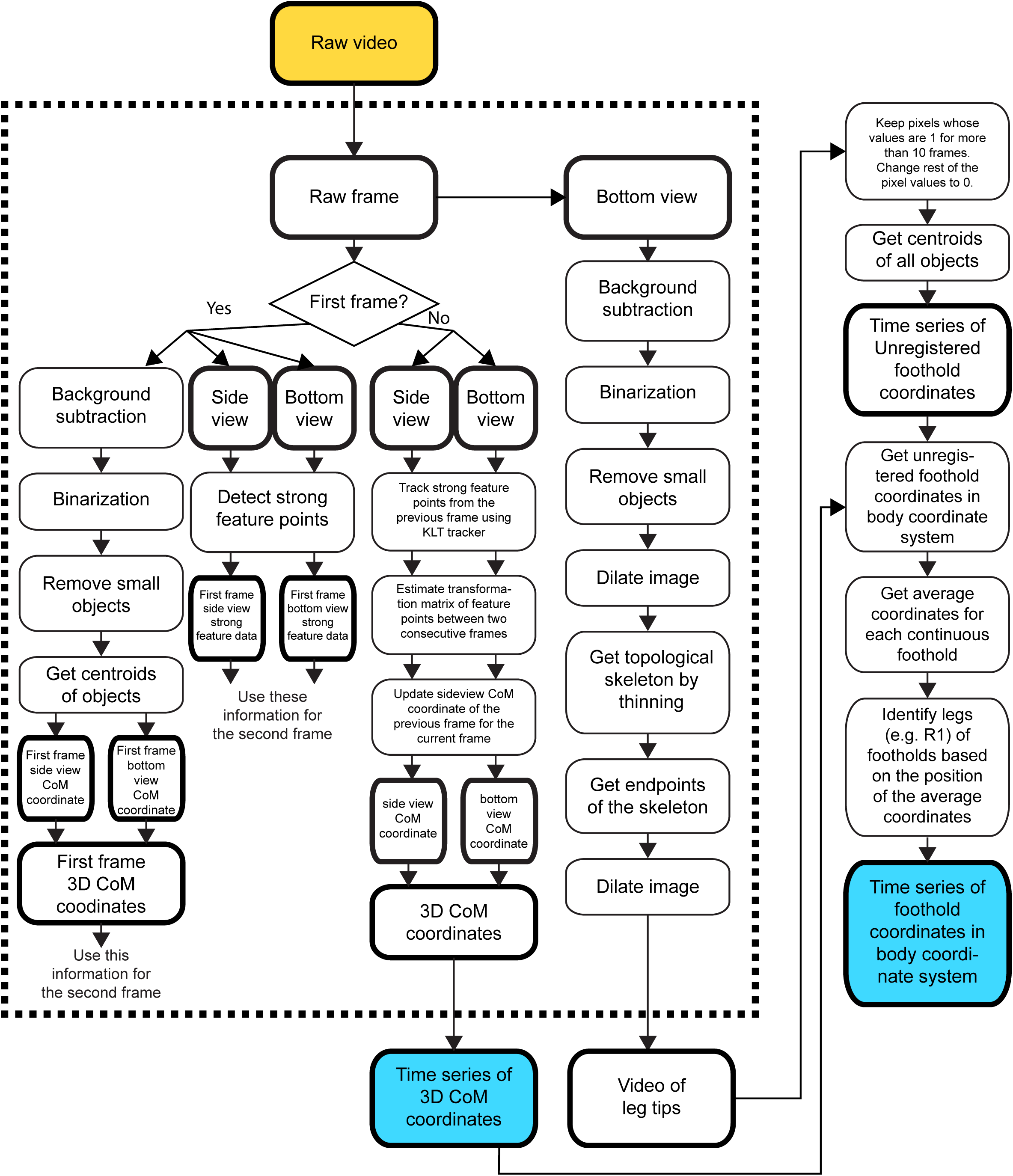
Image processing used to obtain the 3D coordinates of the CoM and the time series of footholds in the body coordinate system. Yellow box denotes input and blue boxes denote outputs. Rectangular elements with thick edges denote processed output. Rectangular elements with thin edges denote processes.

**Supplementary Figure 4.**
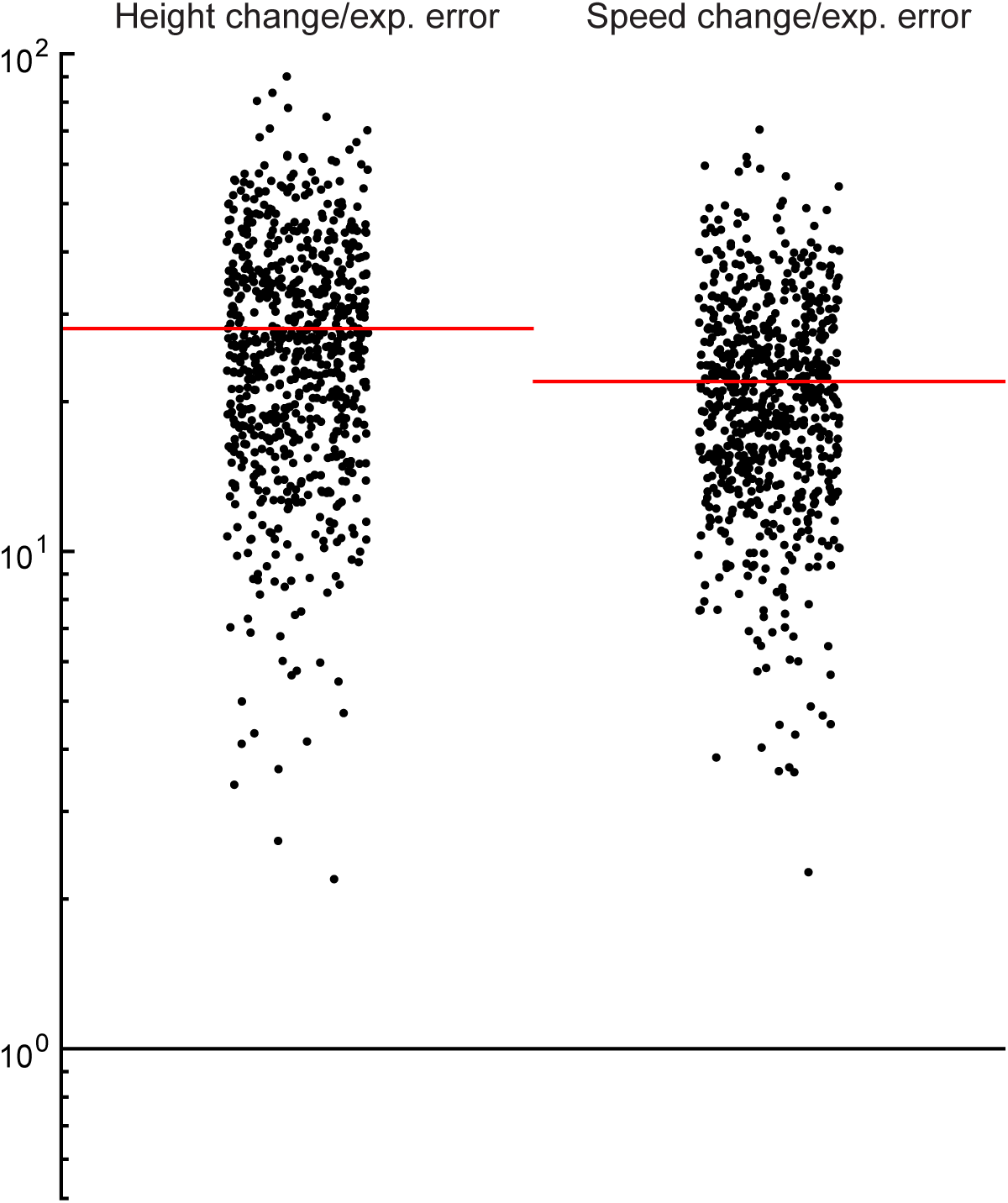
Size of experimental tracking error compared to height change and speed change values. Height change and speed change values were divided by RMSE of tracking. Height change and speed change values are similar but different to the metric defined in Methods 3.3. Here, after detrending a trajectory, we are subtracting minimum from maximum, instead of summing. This quantity indicates the overall amplitude of fluctuation. Red lines are means of the ratios.

**Supplementary Figure 5.**
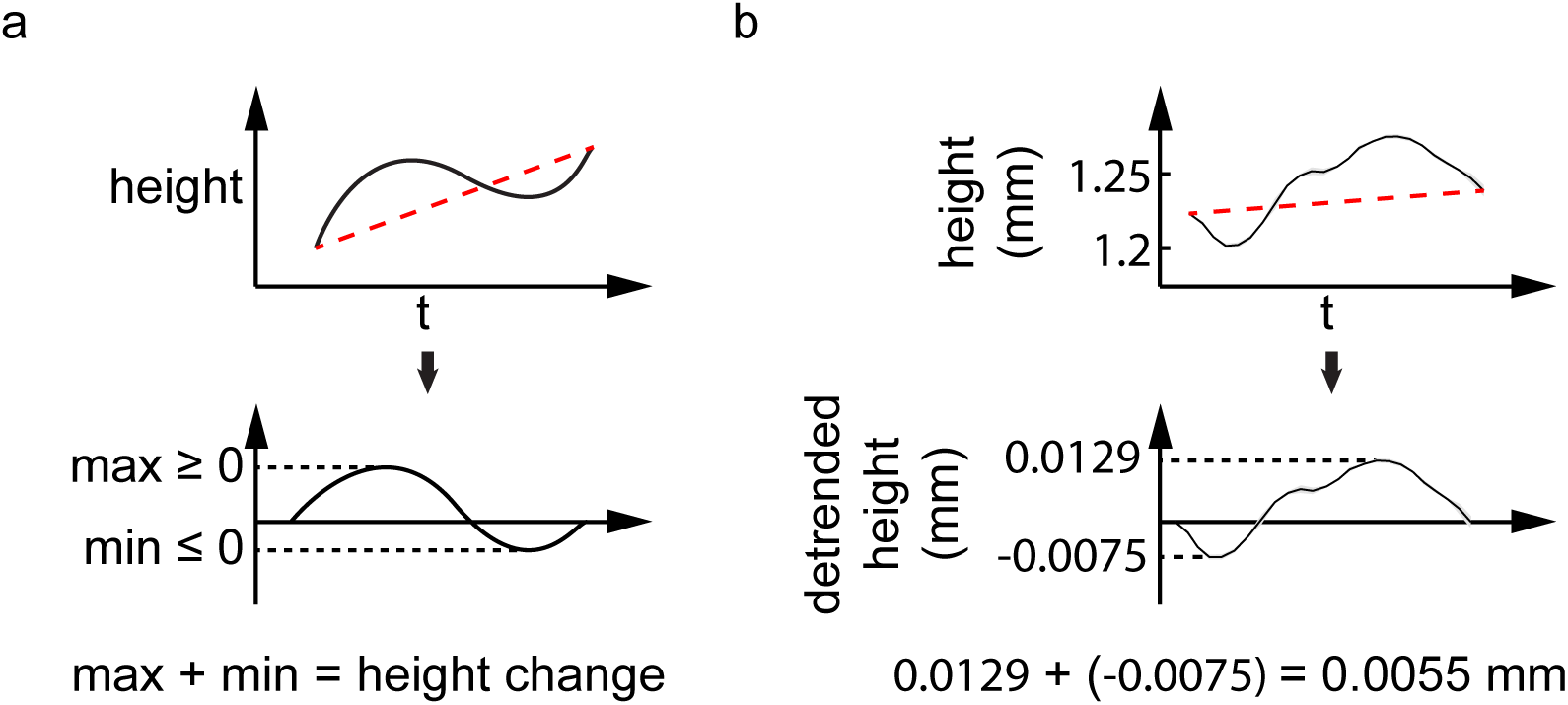
Calculation for height and speed change. (a) Algorithm for height and speed change calculation. (b) Example calculation on the experimental data.

## References

1 Bender, J. A. et al. Kinematic and behavioral evidence for a distinction between trotting and ambling gaits in the cockroach <em>Blaberus discoidalis</em>. The Journal of Experimental Biology 214, 2057-2064, doi: 10.1242/jeb.056481 (2011).

2 Hughes, G. M. The Co-Ordination of insect movements: I The walking movements of insects. Journal of Experimental Biology 29, 267-285 (1952).

3 Wilson, D. M. Insect walking. Annu Rev Entomol 11, 103-122, doi: 10.1146/annurev.en.11.010166.000535 (1966).

4 Wosnitza, A., Bockemuhl, T., Dubbert, M., Scholz, H. & Buschges, A. Inter-leg coordination in the control of walking speed in Drosophila. Journal of Experimental Biology 216, 480-491, doi: 10. 1242/jeb.078139 (2013).

5 Zollikofer, C. Stepping patterns in ants-influence of body morphology. Journal of experimental biology 192, 107-118 (1994).

6 Goldman, D. I., Chen, T. S., Dudek, D. M. & Full, R. J. Dynamics of rapid vertical climbing in cockroaches reveals a template. Journal of Experimental Biology 209, 2990-3000 (2006).

7 Ramdya, P. et al. Climbing favours the tripod gait over alternative faster insect gaits. Nature Communications 8, 14494, doi:10.1038/ncomms14494 (2017).

8 Mendes, C. S., Bartos, I., Akay, T., Marka, S. & Mann, R. S. Quantification of gait parameters in freely walking wild type and sensory deprived Drosophila melanogaster (vol 2, e00231, 2013). Elife 2, doi:UNSP e0056510.7554/eLife.00565 (2013).

9 Strauss, R. & Heisenberg, M. Coordination of Legs during Straight Walking and Turning in Drosophila-Melanogaster. J Comp Physiol A 167, 403-412 (1990).

10 Graham, D. A behavioural analysis of the temporal organisation of walking movements in the 1st instar and adult stick insect (Carausius morosus). Journal of comparative physiology 81, 23-52, doi:10.1007/BF00693548 (1972).

11 Nishii, J. Legged insects select the optimal locomotor pattern based on the energetic cost. Biological cybernetics 83, 435-442 (2000).

12 Smolka, J., Byrne, M. J., Scholtz, C. H. & Dacke, M. A new galloping gait in an insect. Current Biology 23, R913-R915, doi:https://doi.org/10.1016/j.cub.2013.09.031 (2013).

13 Berman, G. J., Choi, D. M., Bialek, W. & Shaevitz, J. W. Mapping the stereotyped behaviour of freely moving fruit flies. Journal of The Royal Society Interface 11, 20140672 (2014).

14 Bässler, U. & Büschges, A. Pattern generation for stick insect walking movements multisensory control of a locomotor program. Brain Research Reviews 27, 65-88 (1998).

15 Schilling, M., Hoinville, T., Schmitz, J. & Cruse, H. Walknet, a bio-inspired controller for hexapod walking. Biological cybernetics 107, 397-419 (2013).

16 Alexander, R. Optimization and gaits in the locomotion of vertebrates. Physiological reviews 69, 1199-1227 (1989).

17 Full, R. J. & Tu, M. S. Mechanics of six-legged runners. Journal of experimental biology 148, 129-146 (1990).

18 Full, R. J. & Tu, M. S. Mechanics of a rapid running insect: two-, four- and six-legged locomotion. Journal of Experimental Biology 156, 215-231 (1991).

19 Blickhan, R. The spring-mass model for running and hopping. Journal of biomechanics 22, 1217-1227 (1989).

20 Blickhan, R. & Full, R. Similarity in multilegged locomotion: bouncing like a monopode. Journal of Comparative Physiology A 173, 509-517 (1993).

21 Cavagna, G. A., Heglund, N. C. & Taylor, C. R. Mechanical work in terrestrial locomotion: two basic mechanisms for minimizing energy expenditure. American Journal of Physiology-Regulatory, Integrative and Comparative Physiology 233, R243-R261 (1977).

22 McMahon, T. A. Muscles, reflexes, and locomotion. (Princeton University Press, 984).

23 McMahon, T. A. & Cheng, G. C. The mechanics of running: how does stiffness couple with speed? Journal of biomechanics 23, 65-78 (1990).

24 Nye, S. W. & Ritzmann, R. E. Motion analysis of leg joints associated with escape turns of the cockroach, Periplaneta americana. Journal of Comparative Physiology A 171, 183-194 (1992).

25 Tomasi, C. & Kanade, T. Detection and tracking of point features. (1991).

26 Morgan, C. L. The Beetle in Motion. Nature 35, 7, doi:10.1038/035007b0 (1886).

27 Pearson, K. The control of walking. Scientific American 235, 72-87 (1976).

28 Biswas, T., Rao, S. & Bhandawat, V. A simple extension of inverted pendulum template to explain features of slow walking✰. Journal of Theoretical Biology 457, 112-123, doi:https://doi.org/10.1016/j.jtbi.2018.08.027 (2018).

29 Couzin-Fuchs, E., Kiemel, T., Gal, O., Ayali, A. & Holmes, P. Intersegmental coupling and recovery from perturbations in freely running cockroaches. J Exp Biol 218, 285-297, doi:10.1242/jeb.112805 (2015).

30 Demoor, J. Recherches sur la marche des Insectes et des Arach-nides. Archiv. Biol 10, 67-608 (1891).

31 Bert, P. Notes diverges sur la locomotion chez plusieurs especes animales. Mint. Soc. Set. Phys. Nat. Bordeaux 4, 59 (1866).

32 Revzen, S. & Guckenheimer, J. M. Estimating the phase of synchronized oscillators. Phys Rev E 78, doi:ARTN 05190710.1103/PhysRevE.78.051907 (2008).

33 Graham, D. Pattern and Control of Walking in Insects. Adv Insect Physiol 18, 31-140 (1985).

34 Gorb, S. N. et al. Structural design and biomechanics of friction-based releasable attachment devices in insects. Integr Comp Biol 42, 1127-1139, doi:DOI 10.1093/icb/42.6.1127 (2002).

35 Bender, J. A. et al. Kinematic and behavioral evidence for a distinction between trotting and ambling gaits in the cockroach Blaberus discoidalis. Journal of Experimental Biology 214, 2057-2064, doi: 10. 1242/jeb.056481 (2011).

36 Bender, J. A., Simpson, E. M. & Ritzmann, R. E. Computer-Assisted 3D Kinematic Analysis of All Leg Joints in Walking Insects. Plos One 5, doi:ARTN e1361710.1371/journal.pone.0013617 (2010).

37 Hoyt, D. F. & Taylor, C. R. Gait and the Energetics of Locomotion in Horses. Nature 292, 239-240 (1981).

38 Alexander, R. M. The Gaits of Bipedal and Quadrupedal Animals. Int J Robot Res 3, 49-59 (1984).

39 Ramdya, P. et al. Climbing favours the tripod gait over alternative faster insect gaits. Nat Commun 8 (2017).

40 Biswas, T., Bhandawat, V. & Rao, S. A new biomechanical template for walking. Integr Comp Biol 57, E206-E206 (2017).

41 Full, R. J. & Koditschek, D. E. Templates and anchors: Neuromechanical hypotheses of legged locomotion on land. Journal of Experimental Biology 202, 3325-3332 (1999).

42 Shi, J. & Tomasi, C. Good features to track. (Cornell University, 1993).

43 Lucas, B. D. & Kanade, T. An iterative image registration technique with an application to stereo vision. (1981).

44 Kalal, Z., Mikolajczyk, K. & Matas, J. in Pattern recognition (ICPR), 2010 20th international conference on. 2756-2759 (IEEE).

45 Buchli, J. & Ijspeert, A. J. Self-organized adaptive legged locomotion in a compliant quadruped robot. Autonomous Robots 25, 331 (2008).

46 Wilshin, S. et al. Longitudinal quasi-static stability predicts changes in dog gait on rough terrain. Journal of Experimental Biology, jeb. 149112 (2017).

47 Marple, L. Computing the discrete-time” analytic” signal via FFT. IEEE Transactions on signal processing 47, 2600-2603 (1999).

48 Boashash, B. Estimating and interpreting the instantaneous frequency of a signal. II. Algorithms and applications. Proceedings of the IEEE 80, 540-568 (1992).

49 Smith, J. O. Mathematics of the Discrete Fourier Transform (DFT) with Audio Applications, Second Edition. W3K Publishing (2007).

50 Dixon, L. C. W. The Global Optimization Problem. An Introduction. Toward global optimization 2, 1-15 (1978).

